# Not1 and Not4 inversely determine mRNA solubility that sets the dynamics of co-translational events

**DOI:** 10.1101/2022.03.14.484207

**Authors:** George Allen, Benjamin Weiss, Olesya Panasenko, Susanne Huch, Zoltan Villanyi, Benjamin Albert, Daniel Dilg, Marina Zagatti, Paul Schaughency, Susan E. Liao, Jeff Corden, Christine Polte, David Shore, Zoya Ignatova, Vicent Pelechano, Martine A. Collart

**Author notes:** Department of Biomolecular Sciences, The Weizmann Institute of Science, 76100 Rehovot, Israel (Benjamin Weiss). Molecular, Cellular & Developmental Biology (MCD), Center for Integrative Biology (CBI), University of 11 Toulouse CNRS/UPS, Bâtiment IBCG, 118, route de Narbonne, 31062 Toulouse cedex 9, France. NYU, Courant Institute of Mathematical Sciences, New York, USA. Equal contributing first authors (George Allen, Benjamin Weiss).

## Abstract

**Background:** The Ccr4-Not complex is most well known as the major eukaryotic deadenylase. However, several studies have uncovered roles of the complex, in particular of the Not subunits, unrelated to deadenylation and relevant for translation. In particular, the existence of Not condensates that regulate translation elongation dynamics have been reported. Typical studies that evaluate translation efficiency rely on soluble extracts obtained after disruption of cells and ribosome profiling. Yet cellular mRNAs in condensates can be actively translated and may not be present in such extracts.

**Results:** In this work, by analyzing soluble and insoluble mRNA decay intermediates in yeast, we determine that insoluble mRNAs are enriched for ribosomes dwelling at non-optimal codons compared to soluble mRNAs. mRNA decay is higher for soluble RNAs, but the proportion of co-translational degradation relative to the overall mRNA decay is higher for insoluble mRNAs. We show that depletion of Not1 and Not4 inversely impact mRNA solubilities and, for soluble mRNAs, ribosome dwelling according to codon optimality. Depletion of Not4 solubilizes mRNAs with lower non-optimal codon content and higher expression that are rendered insoluble by Not1 depletion. By contrast, depletion of Not1 solubilizes mitochondrial mRNAs, which are rendered insoluble upon Not4 depletion.

**Conclusion:** Our results reveal that mRNA solubility defines dynamics of co-translation events and is oppositely regulated by Not1 and Not4, a mechanism that we additionally determine may already be set by Not1 promoter association in the nucleus.

## Introduction

Adequate regulation of gene expression is essential for health, fitness and development of all living organisms. While transcription is the most immediate and focal point of gene regulation, gene expression is also importantly controlled at post-transcriptional levels (1–4). Repression of translation initiation, the major rate-limiting step of translation, for instance, plays a key role in cellular responses to nutrient levels and stresses (5–7). Nevertheless, translation output can also be regulated at the elongation step, according to the availability of charged tRNAs, codon bias, the amino acid composition of the nascent chain, co-translational folding, interactions of nascent chains with auxiliary factors, and by mRNA localization or mRNA partitioning into membrane-less granules (8–14). mRNA-protein condensates were first associated with translational repression (stress granules) and mRNA decay (p-bodies) (15), but recent evidence indicates active translation in stress granules (16), and positive roles of granules for translation have been proposed (12, 17–19). The development of techniques such as ribosome profiling (Ribo-Seq) (20), visualizing with codon-specific precision the position of ribosomes on mRNAs genome-wide, or the sequencing of 5’P decay intermediates (5’P-Seq) (21), revealing patterns of co-translational decay intermediates, has enabled the analysis of translation elongation dynamics with unprecedented depth and precision.

The conserved Ccr4-Not complex plays a key role in mRNA metabolism (22). First identified as a transcriptional regulator (23–25), a role later confirmed (26–30), Ccr4-Not is most well-known as the major eukaryotic deadenylase (31–33). Thereby it is central in mRNA turnover and translational repression (34, 35). It is generally active for post-translational mRNA decay, but can also be tethered to mRNAs by RNA binding proteins or the microRNA machinery (36–39). Ccr4-Not can also inhibit translation independently of deadenylation (40) or activate decapping (41).

Ccr4-Not additionally regulates translation and co-translational processes. Ribosome-associated proteins, notably the nascent polypeptide ribosome associated complex (NAC) and Rps7A (42, 43) are known targets of Not4 ubiquitination. The ubiquitination-deubiquitination cycles of Rps7A are important for translation (44) and non-ubiquitinated Rps7A enables translation elongation through polyarginine stretches that normally provoke ribosome stalling (17). Rps7A ubiquitination is also important for translation regulation during ER stress (45). Not proteins co-sediment with polysomes (42, 46) and Not5 polysome-association is promoted by Rps7A ubiquitination. Furthermore, the co-translational association of proteins is impaired in the absence of Not4 or Not5 (47–49) and co-localization of mRNAs encoding two subunits of the proteasome that assemble co-translationally depends upon Not1 (18). It was recently shown that Not5 associates with the ribosomal E site in post-translocation state providing thereby a means for the Ccr4-Not complex to monitor the translating ribosome according to codon optimality (50). It was proposed that this regulates the turnover of mRNAs, consistent with Not5-dependent longer half-lives of mRNAs with a high content of non-optimal codons (50). We recently proposed an alternative role for Not4 and Not5, consistent with Not5 monitoring the translating ribosome according to codon optimality (17). In our model we proposed that Not5 can tether ribosome-nascent chain complexes (RNCs) to condensates that exclude the translation initiation and elongation factor eIF5A (51, 52). A central role of Not4 and Not5 in translation elongation dynamics is corroborated by the fact that, when deleted in cells, newly synthesized proteins massively aggregate (42, 53) accompanied with a high level of abortive translation products (17).

Condensate mRNAs, like other insoluble mRNAs such as membrane-associated mRNAs, are not captured by typical ribosome and polysome profiling approaches. In this work we used the sequencing of mRNA decay intermediates to address whether soluble mRNAs and insoluble mRNAs have different translation elongation dynamics and if different mRNA classes partition differently into soluble and insoluble mRNA pools. We further investigated if elongation dynamics of the different mRNA pools was altered immediately upon depletion of Not1, Not4 and Not5. Taken together, our data indicate that ribosomes dwell at non-optimal codons in the non-soluble RNA fraction. In turn, soluble mRNAs show more mRNA 5’ to 3’ degradation, but proportionally less co-translational decay. Additionally, we determine that the depletion of Not1 and Not4 regulate mRNA partitioning between soluble and insoluble fractions in an opposing manner, which in turn correlates with the opposite impacts on ribosome dwelling at optimal and non-optimal codons in the soluble RNA pool. Our results are compatible with a model whereby mRNA solubility sets the dynamics of co-translational events and is regulated by the Not proteins.

## Results

### Paused ribosomes at non-optimal codons are enriched within non-soluble mRNA fractions

In a recent study we showed that the dynamics of translation elongation were greatly affected in the absence of Not4 and Not5 in *S.cerevisiae* and we proposed that this was due to defective tethering of translating ribosomes to ribonucleoprotein (RNP) condensates, in particular ribosomes paused at non-optimal codons (17). This model predicts that ribosomes paused at non-optimal codons should be enriched in the insoluble mRNA condensates. To test this idea, we needed a means to detect insoluble mRNAs. To achieve this, we compared the cell’s total mRNA pool that can be prepared from cell pellets (called hereafter “total RNA”) and includes insoluble mRNAs, and the cell’s soluble mRNA pool obtained from cell lysates (called hereafter “soluble RNA”). We investigated the distribution of mRNAs between the soluble and total RNA pools (referred to from here onwards as “solubility”) and the 5’P mRNA decay intermediates in total versus soluble RNA pools. We grew wild-type cells in rich medium to exponential phase and split them in two, extracting the total RNA from one aliquot and the soluble RNA from the other, in biological triplicates. For normalization we spiked in each sample a constant amount of RNA from *S. pombe*. For each sample one aliquot was subjected to RNA-Seq to determine the transcriptome and the other to 5’P-Seq to determine mRNA decay intermediates that can provide information on co-translational decay and ribosome dwelling (21)(**Table S1**).

We first determined overall mRNA solubilities (log2FC soluble/total). mRNA levels within the soluble and total mRNA pools correlated overall (**Figure 1A**). Nevertheless, the mRNA solubilities spanned a relatively wide range, from −2.744 to 1.791 (**Figure 1B**). It is interesting to note that a GO-term analysis of the mRNAs showing the lowest solubilities revealed “endoplasmic reticulum”, “membrane” and “cell wall” (**Figure S1A**). Membranes are expected to sediment in the first centrifugation step after cell lysis, and mRNAs encoding membrane proteins or proteins that must transit through membranes can be targeted to membranes during translation (12, 54–59). Hence, such mRNAs can indeed be expected to be depleted from soluble extracts.

**Figure 1.**
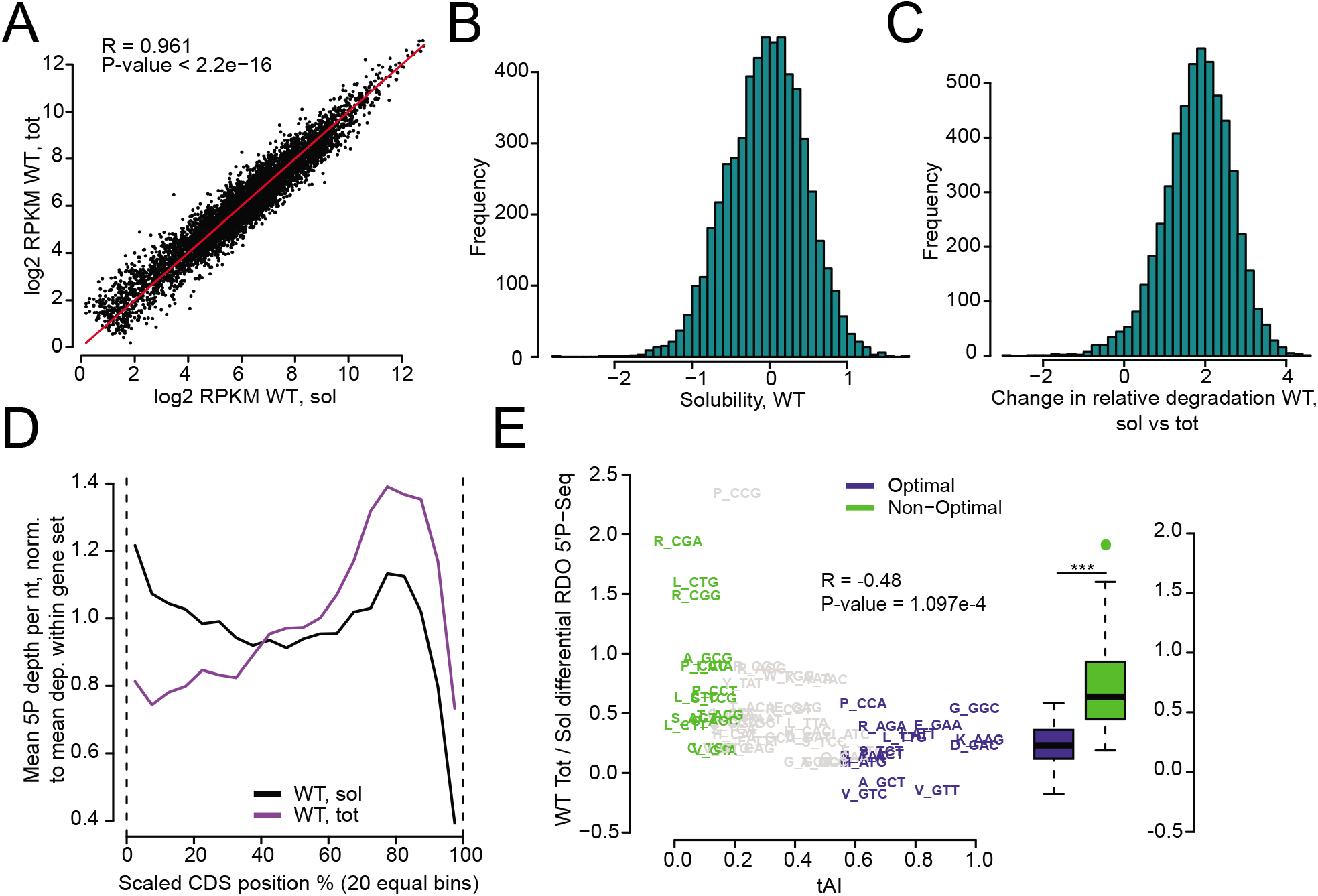
mRNAs that are less soluble are enriched in non-optimal codons. **A**. Scatterplot comparing RPKMs of mRNAs in soluble (sol) and total (tot) mRNA pools. **B**. Distribution of solubility of mRNAs, defined as log2FC soluble/total from DESeq2 in RNA-Seq. **C**. Distribution of relative degradation levels of 5’P mRNA intermediates compared to the RNA abundance (log2FC 5’P-Seq/RNA-Seq from DESeq2 using spike-in) in soluble versus total RNA pools. **D**. Metagene profile dividing each CDS into 20 equal bins and finding the mean normalized reads of 5’P mRNA intermediates in each for soluble and total RNA pools in wild type cells. **E**. Scatterplot comparing differential 5’P-RDOs in total versus soluble RNA pools with the tRNA adaptation index (tAI). The 15 most optimal codons are indicated in blue, the 15 most non-optimal codons are shown in green. The optimal and non-optimal codon relative RDOs were compared with a one-sided Wilcoxon rank-sum test giving a p-value of 5.627e-05.

To investigate the distribution of 5’P mRNA decay intermediates between soluble and total RNA fractions we compared the level of mRNA degradation intermediates determined with 5’P-Seq to the total mRNA level defined with RNA-Seq (**Table S1**). We used the spike-ins to normalize 5’P-Seq reads to the total RNA-Seq reads. This comparison provides a snapshot of the fraction of mRNAs undergoing degradation and we refer to this measure as “relative degradation” from here onwards. The relative degradation was higher in the soluble mRNA pool than in the total RNA pool, as indicated by the shift of distribution between the two pools centered around 2 (**Figure 1C**). In addition, metagene profiles of 5’P-Seq depth (i.e. 5’P reads normalized to library size) across coding sequences (CDSs) were significantly different between the soluble and total RNA pools (**Figure 1D**). For the soluble RNA pool, we noted high levels of 5’P mRNA ends mapping throughout the CDS, while for the total RNA the 5’P reads were lower at the beginning of the CDS and higher at the end. These metagene profiles were similar for mRNAs with high, medium or low amount of 5’P reads. The exception were shorter mRNAs that showed no accumulation of 5’P reads at the end of the CDSs (**Figure S1B**). For both RNA pools, there was a drop in 5’P reads within the last 150 nucleotides of CDSs, despite our use of a mix of random hexamer and oligo(dT) priming for the library preparation facilitating recovery of regions proximal to the poly(A) site (60). This is due to the fact that we look at the 5’ region of libraries of a specific insert size. The metagene profile of 5’P decay intermediates for soluble mRNAs was similar to the metagene profile of ribosome footprints observed previously (**Figure S1C**) (17).

The difference in metagene profiles of 5’P decay intermediates from soluble and total RNAs could be due to differences in the processivity of 5’ to 3’ decay in the soluble and total mRNA pools, directly or indirectly linked to differences in velocities of ribosomes. Ribosome profiling data provides information on ribosome footprints for the soluble mRNA pool. No similar data can be obtained for insoluble RNAs. However, 5’P-Seq data is informative on ribosome dwelling (21). Indeed, in the case of co-translational mRNA decay, the progression of the 5’ to 3’ Xrn1 exonuclease can be limited by the dwelling of the last translating ribosome, depending upon relative kinetics of decay and ribosome progression, and the onset of decay compared to progression of the last translating ribosome. Thus, we used 5’P-Seq to compare ribosome dwelling in total and soluble RNA pools. We defined A-site ribosome dwelling occupancies from the 5’P-Seq data (5’P-RDOs), using the 5’P reads 17 nucleotides upstream of each codon. The differential 5’P-RDOs between the total and soluble RNA pools anticorrelated with codon optimality (**Figure 1E**), indicating that co-translational decay intermediates accumulating at ribosomes paused on non-optimal codons were enriched in total RNAs compared to soluble RNAs. Since the total RNA pool includes both soluble and insoluble RNAs, these results suggest that ribosome dwelling at non-optimal codons in their A-site is higher in non-soluble RNA fractions.

mRNA turnover can occur by mechanisms other than co-translational degradation. To get some idea of this for both RNA pools we generated metagene profiles of mRNA 5’ ends generated by RNA-Seq and compared them to the metagene profiles of 5’P-Seq. The metagene profiles of 5’end reads of the RNA-Seq were more similar between soluble and total RNA pools (**Figure S1D**) than the 5’P-Seq metagene profiles (**Figure 1D**), particularly there was less difference at the 5’ end of CDSs. This was also apparent by evaluating for soluble and total RNAs reads in the second half of the CDSs (70-90%) to those in the first half (10-30%) for 5’P-Seq and for the 5’end of the RNA-Seq reads (**Figure S1E**).

For soluble and total RNAs, the 5’P-RDO changes in the second half of the CDS compared to the first half were inversely correlated with codon optimality (**Figure S1F**). These observations could result from the last translating ribosome dwelling longer at non-optimal codons in the second half compared to the first half of CDSs. Alternatively, they could be explained by a delayed decay onset after the last translating ribosome, a model that seems more likely considering that longer mRNAs show more 5’P decay intermediates at the end of CDSs (see above, **Figure S1B**).

### Solubility of mRNAs is inversely modified upon depletion of Not1 or Not4

We next investigated the direct role of Not proteins in regulating mRNA solubilities and differences in 5’P-RDOs between total and soluble RNA pools. For this, we created Not4 and Not5 auxin-inducible degron strains, along with the Not1 degron strain described previously (17). Expression of the degron-regulated proteins was abolished after 15 min of auxin treatment (**Figure S2A**). Expression of Not4 was not altered after depletion of Not5, and Not5 expression was not altered after depletion of Not4, and in both strains, expression of Not1 was unaffected.

Libraries from soluble and total RNA pools were generated from the degron strains and their isogenic wild-type counterpart following 15 min of auxin treatment (**Table S1**). As shown above, the expression of mRNAs in total and soluble RNA pools correlated for wild-type cells. However, we noted that the correlation was lower upon Not1 and Not4 depletion, and minimally affected upon Not5 depletion (**Figure 2A**). Some mRNAs showed much less solubility upon Not1 depletion and inversely some mRNAs showed much more solubility upon Not4 depletion (**Figure 2B**). Interestingly, there was an overall inverse correlation between the changes in mRNA solubility upon Not1 and Not4 depletion (**Figure 2C**). The depletion of Not5 exhibited marginal effects on the mRNA solubility and the subtle changes somewhat resembled those observed upon Not4 depletion (**Figure S2B**, left panel) but not those observed upon Not1 depletion (**Figure S2B**, right panel).

**Figure 2.**
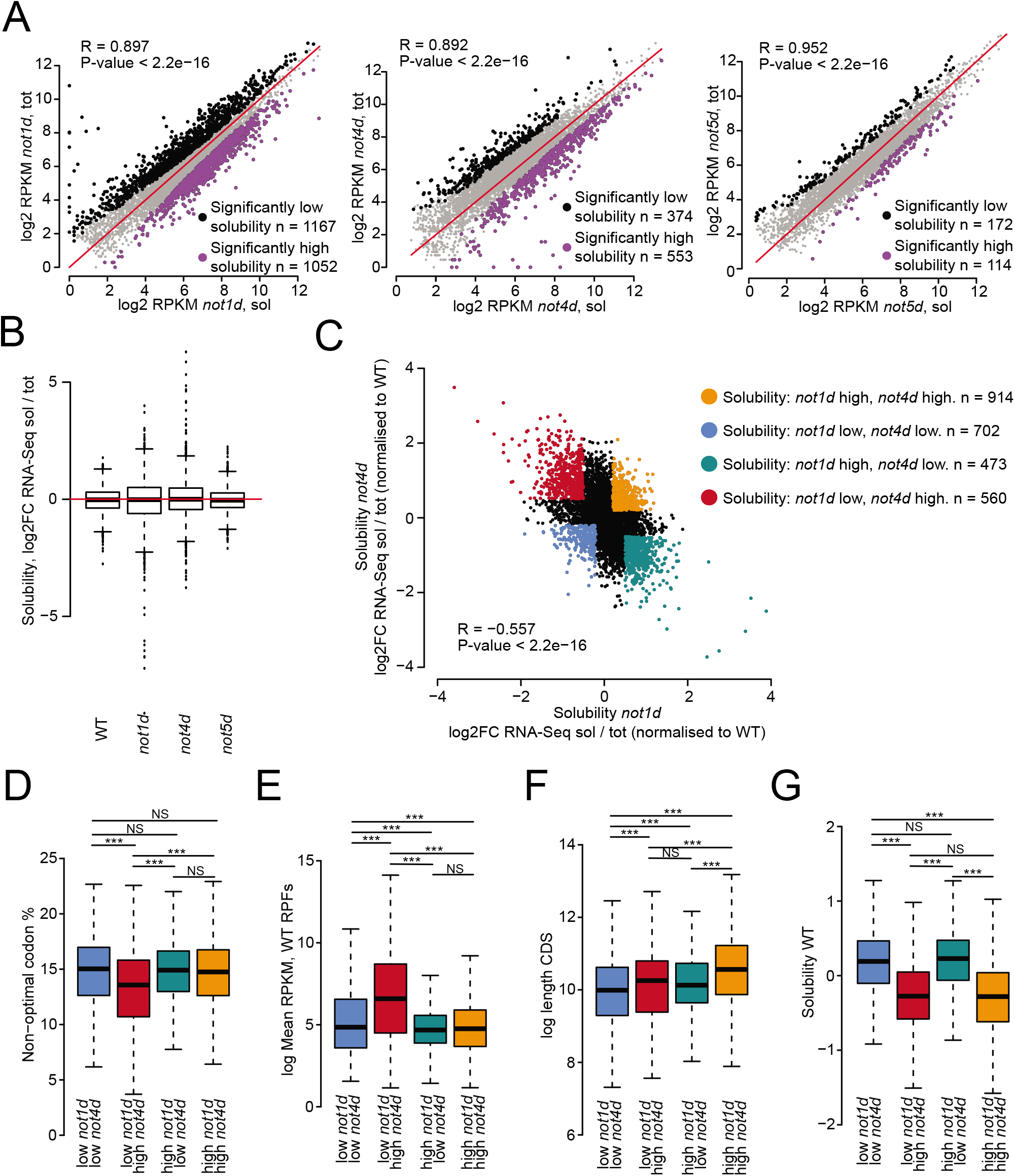
Not1 and Not4 inversely regulate mRNA solubilities. **A**. Scatterplot comparing RPKMs of mRNAs in soluble (sol) and total (tot) mRNA pools before and after Not1 (*not1d*), Not4 (*not4d*) and Not5 (*not5d*) depletion (from left to right) in soluble and total mRNA pools. mRNAs significantly less soluble after depletion are indicated in black and more soluble in purple (cutoffs from DESeq2 RNA-Seq sol/tot - high solubility: [log2FC > 0, FDR < 0.05] OR [log2FC > 1, p-value < 0.05]; low solubility: [log2FC < 0, FDR < 0.05] OR [log2FC < −1, p-value < 0.05]). **B**. Box plot analysis indicating mRNA solubilities in cells before (WT) or after Not1, Not4 and Not5 depletion. **C.** Scatterplot comparing changes in mRNA solubilities before and after Not1 and Not4 depletion. **D-G**. Box plot analysis comparing features of mRNAs falling into the four categories defined by changes in mRNA solubilities upon Not1 and Not4 depletion color coded in panel **C** with regard to **D**: content in non-optimal codons, **E**: abundance (RPF RPKMs), **F**: length and **G**: solubility in wild type cells. Number of mRNAs in box plots, left to right: 823, 637, 522 and 1040. Significance of differences is indicated at the top of the box plots – p-values are calculated using a two-sided Welch two sample t-test.

We focused our attention on the mRNA sets that showed opposing changes in solubility following Not1 and Not4 depletion. This concerned 1158 mRNAs (**Table S2**), of which 637 were less soluble upon Not1 depletion but more soluble upon Not4 depletion (solubility not1d/WT < −0.5 and not4d/WT > 0.5, red category on **Figure 2C**) and instead 522 mRNAs more soluble upon Not1 depletion and less soluble upon Not4 depletion (solubility not1d/WT > 0.5 and not4d/WT < −0.5, green category on **Figure 2C**). We checked for specific features of these mRNAs such as their expression level, length and content in non-optimal codons, and solubility in wild type cells (**Figure 2D–G**). mRNAs with lowered solubility upon Not1 depletion and instead higher solubility upon Not4 depletion (red category) had a significant lower content in non-optimal codons (**Figure 2D**) and were higher expressed (**Figure 2E**). mRNAs with higher solubility upon Not1 depletion and instead lower solubility upon Not4 depletion (green category) were enriched for the GO-term “mitochondrial organization” and “mitochondrial translation” (**Figure S2C**). mRNAs with reduced solubility upon either Not1 or Not4 depletion (blue category) were distinguishable from those with higher solubility (orange category) by their shorter length (**Figure 2F**) and were enriched for the GO-term “endocytosis”, whilst the mRNAs with higher solubility upon either Not1 or Not4 depletion (orange category) were enriched for the GO-term “RNA modification” and “tRNA processing” (**Figure S2C**). Notably, mRNAs that were more soluble upon Not4 depletion were mRNAs that tended to be less soluble in wild type cells (**Figure 2G**).

Taken together, these results indicate that specific features of the mRNAs, or specific mRNA families, indicates how their solubility will be impacted upon depletion of Not1 or Not4.

### Following depletion of the Not proteins changes in solubility and relative degradation correlate

We next determined how the relative degradation of the total and soluble RNA pools was impacted upon depletion of the Not proteins by comparing the fraction of molecules undergoing degradation to the total mRNA abundance. The overall higher relative degradation in the soluble RNA pool compared to the total RNA pool was maintained following depletion of the Not proteins (**Figure 3A**). However, the relative degradation of the soluble mRNA pool was decreased upon Not1 depletion but increased upon Not4 depletion, and to a lesser extent upon Not5 depletion. For the total RNA pool, the relative degradation slightly increased following depletion of any of the Not proteins.

**Figure 3.**
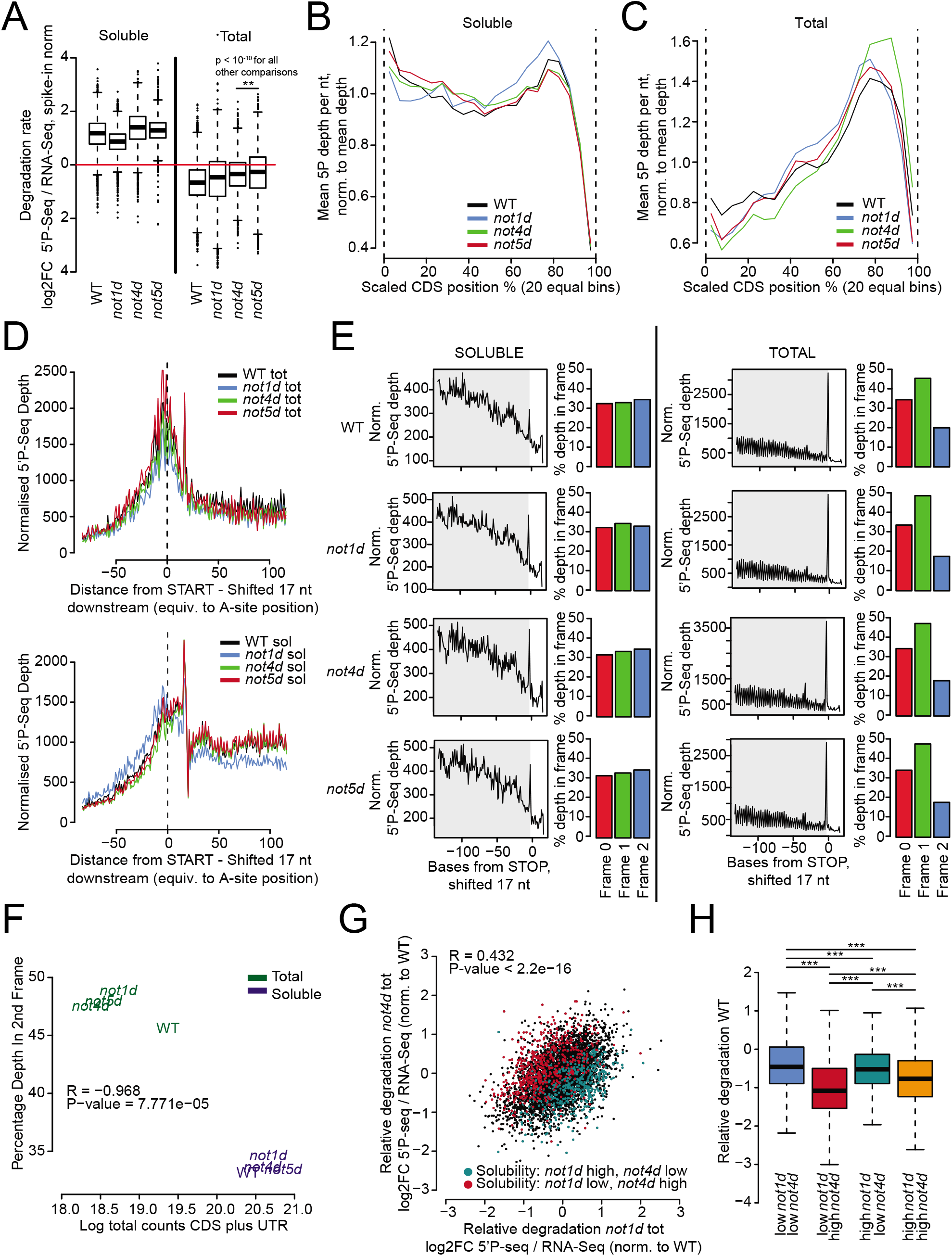
Soluble mRNAs show more relative degradation but less detectable co-translational decay than insoluble mRNAs. **A.** Box plot analysis indicating levels of 5’P mRNA decay intermediates compared to total RNA normalized by spike in control RNA in cells before (WT) and after Not1 (*not1d*), Not4 (*not4d*) and Not5 (*not5d*) depletion, in soluble and total RNA pools. Significance of differences is indicated at the top of the box plots - p-values are calculated using a two-sided Welch two sample t-test. **B-E**. Positions of 5’ ends of decay intermediates were shifted 17nt downstream to simulate the A-site of the adjacent ribosome and create metagene profiles of 5’P mRNA decay intermediates for WT and after Not1, Not4 and Not5 depletion; **B** and **C**: over the whole CDS for soluble (**B**) and total (**C**) RNA pools; **D**: around the start codon for the total (upper panel) or the soluble (lower panel) RNA pools; **E**: before the stop codon in soluble (left) and total (right) RNA pools, with an indication of the percentage of 5’P decay intermediate 5’ ends (no shift) in each of the reading frames over the entire ORFs for the cells before (WT, first row) and after Not1 (*not1d*, second row), Not4 (*not4d*, third row) and Not5 (*not5d*, fourth row) depletion. **F**. Scatterplot representing percentage of 5’P decay intermediate 5’ ends in Frame 1 within the soluble and total RNA pools for the indicated strains. **G.** Scatterplot comparing relative degradation (corrected to WT) in *not1d* tot and *not4d* total RNAs. Transcripts with solubility high in *not1d* and low in *not4d* are indicated in green, low in *not1d* and high in *not4d* are red, related to **Figure 2C**. **H**. Same analysis as **Figure 2D** but for relative degradation in wild type cells.

The metagene profiles of 5’P decay intermediates for the soluble mRNAs were overall similar before and after Not protein depletion, except for a slight decrease in 5’P decay intermediates in the first half of the CDS and increase in the second half of the CDS upon Not1 depletion (**Figure 3B**). For the total RNAs, a decrease in 5’P mRNA ends in the first half of the CDS and an increase in the second half of the CDS was observed upon depletion of each of the Not proteins, most prominently upon Not4 depletion (**Figure 3C**).

We looked more closely in the region surrounding the beginning of the CDS. For the total RNA pool, there was a peak of 5’P mRNA ends centered around the start codon (**Figure 3D**, upper panel). For the soluble RNAs, a peak at start was so not well pronounced (**Figure 3D**, lower panel). In both soluble and total RNAs a peak at about 20 nucleotides downstream of the start codon was detectable. Notably, a peak of ribosome footprints around this position has been seen in ribosome profiling data sets (eg (61)). The metagene profiles around the start were similar before and after depletion of the Not proteins, with the exception of the slightly increased reads upstream of the start and decreased reads downstream of start for the soluble mRNAs upon Not1 depletion that could be indicative of more ribosome pausing at start.

We next focused on the region before the stop codon. In particular, we inspected the profiles for three nucleotide-periodicity expected for 5’P mRNA reads resulting from co-translational decay. Periodicity was well detectable for the total RNAs but not for the soluble RNAs (**Figure 3E**). This three nucleotide-periodicity was improved for the total RNA pool upon depletion of each of the Not proteins (**Figure 3F**), and this was unrelated to the depth of the sequencing libraries (**Figure S2D**). For the total RNA samples, but not for the soluble RNAs, an important peak was detected at the stop codon, suggesting that ribosome recycling might be slow for insoluble mRNAs. Upon depletion of each Not protein, the peak at the stop codon increased in the soluble RNAs and it increased for the total RNAs upon Not4 depletion.

The pool of mRNAs that showed the most extreme change in solubilities upon Not1 and Not4 depletions (**Figure 2C**) were distinguishable by opposing behaviors with regard to their relative degradation: those mRNAs more soluble upon Not1 or Not4 depletion tended to have increased relative degradation (**Figure 3G**). We also noted that mRNAs that become more soluble upon Not4 depletion and tended to be less soluble in wild type cells as mentioned above, consistently also had less relative degradation in wild type cells (**Figure 3H**). These findings indicate that changes in solubility and relative degradation correlate.

### Metagene profiles of decay intermediates distinguish mRNAs inversely impacted by Not1 and Not4

We focused on the mRNAs that showed opposite solubility regulations upon Not1 and Not4 depletion (green and red categories of **Figure 2C**). The 5’P-Seq metagene profiles for these 2 groups of mRNAs were very different (**Figure 4A** and see box plots on **Figure 4B** for quantification of reads in the total versus soluble RNA pools for different portions of the CDS). This was most striking for the soluble RNA pool, whereby for the mRNAs of the red category (less soluble upon Not1 depletion) there was a constant lower amount of reads in the first half of the CDS and increased reads in the second half of the CDS. Instead, for the mRNAs of the green category (more soluble upon Not1 depletion) the reads decreased in the second half of the CDS and showed a peak of reads centered at 20% of the CDS. For this latter category the reads for the total RNA pool were less different between the first and second halves of the CDS than for the first category that showed a pattern more similar to that seen for all mRNAs.

**Figure 4.**
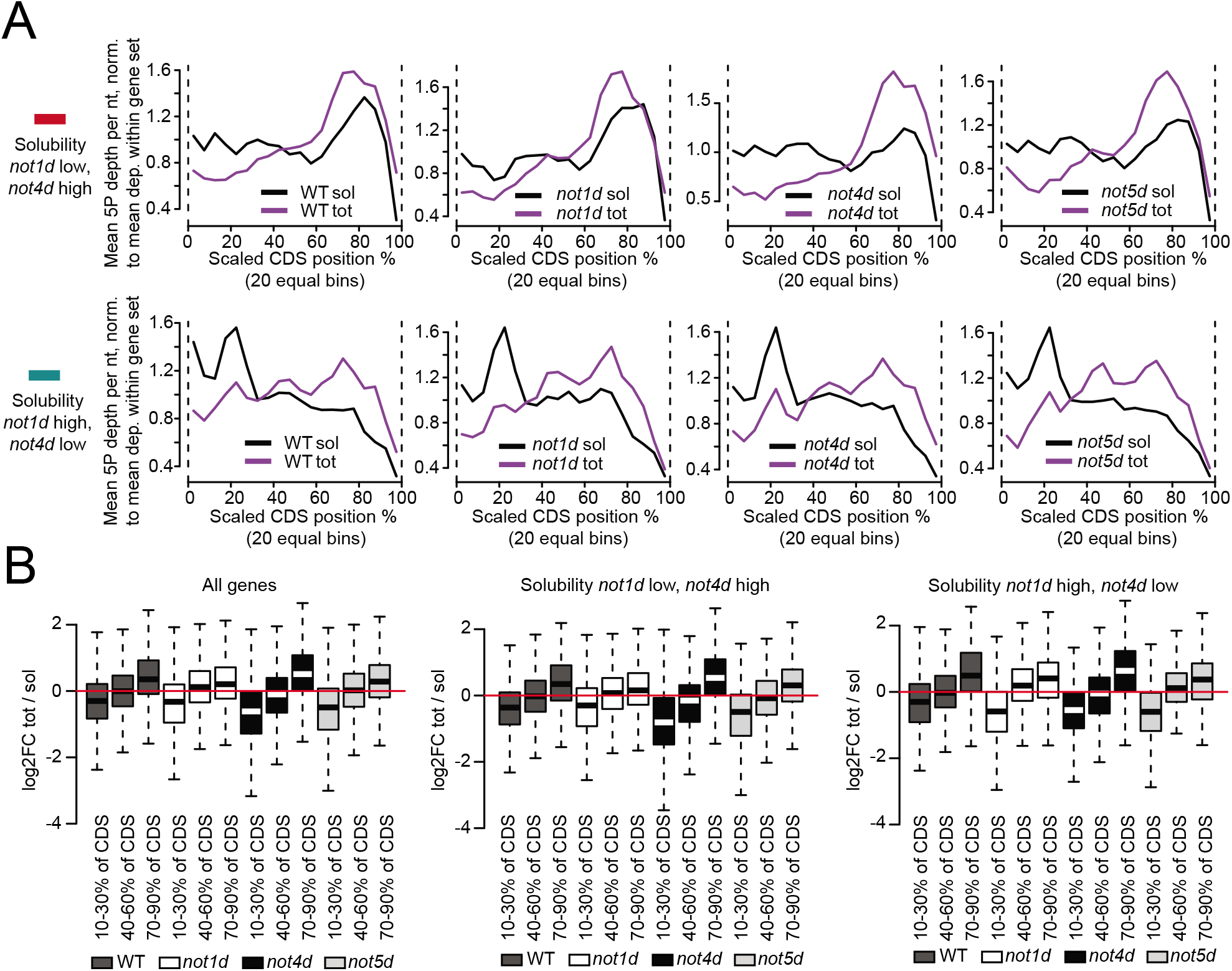
mRNAs with solubilities inversely regulated by Not1 and Not4 show very different co-translational decay patterns. **A**. Metagene analysis of 5’P-Seq for total and soluble mRNAs of the red and green categories of **Figure 2C** in wild type and after Not1, Not4 or Not5 depletions. **B**. Box plot analysis of the 5’P-Seq reads for total versus soluble mRNAs, comparing proportion of reads falling between 10-30%, 40-60% and 70-90% of CDSs, for all mRNAs (left), mRNAs of the red category (middle) or green category (right). Transcripts are only included in the analysis if their CDS is covered by at least 20 5’P-Seq reads in both soluble and total.

For mRNAs less soluble upon Not1 depletion (red mRNAs) (**Figure 4A**, upper panels and **Figure 4B**, middle panel), depletions of Not4 or Not5 decreased 5’P-Seq reads in the total versus soluble RNA pools in the first third (10-30%), whereas Not1 depletion slightly increased 5’P-Seq reads in the total versus soluble RNA pools in the first third (10-30%) and decreased them in the last third (70-90%) of the CDS. For the mRNAs less soluble upon Not4 depletion (green mRNAs) (**Figure 4A**, lower panels and **Figure 4B**, right panel), depletions of Not1 or Not5 increased reads in the middle of the CDS (40-70%) for the total RNAs.

These results show that co-translational decay patterns are very different for the two mRNA categories whose solubility is inversely impacted by Not1 or Not4 depletion. They additionally suggest that Not5 works with Not4 for the red mRNA category but with Not1 for the green category.

### A-site RDOs calculated from 5’P Seq data distinguish soluble and total RNA pools and the actions of the different Not proteins

We compared A-site 5’P-RDO changes of the soluble RNA pools following Not1, Not4 or Not5 depletions. In the soluble RNA pool, the A-site 5’P-RDO changes observed upon Not4 and Not5 depletion correlated and they correlated with codon optimality (**Figure 5A**, left panel). Instead, the A-site 5’P-RDO changes observed upon Not1 and Not5 depletion showed a minor anti-correlation (**Figure 5A**, right panel). Such an inverse correlation was more pronounced and significant for the A-site 5’P-RDO changes observed upon Not1 and Not4 depletion (**Figure 5B**, left panel). It was abolished if the mRNAs of the red and green categories were removed (**Figure 5B**, middle panel) and was instead more important and much more significant when only the mRNAs of the red and green categories were analyzed (**Figure 5B**, right panel), with a clear codon-optimality related effect. It is interesting to note that RDO changes in cells lacking Not5 compared to wild type cells measured by Ribo-Seq previously and shown to correlate with 5’P-RDO changes upon Not1 depletion (17) showed instead a mild inverse correlation with 5’P-RDO changes upon Not5 depletion (**Figure S3A**). This suggests that in *not5Δ* cells limiting amounts of Not1 have a dominant impact on RDOs.

**Figure 5.**
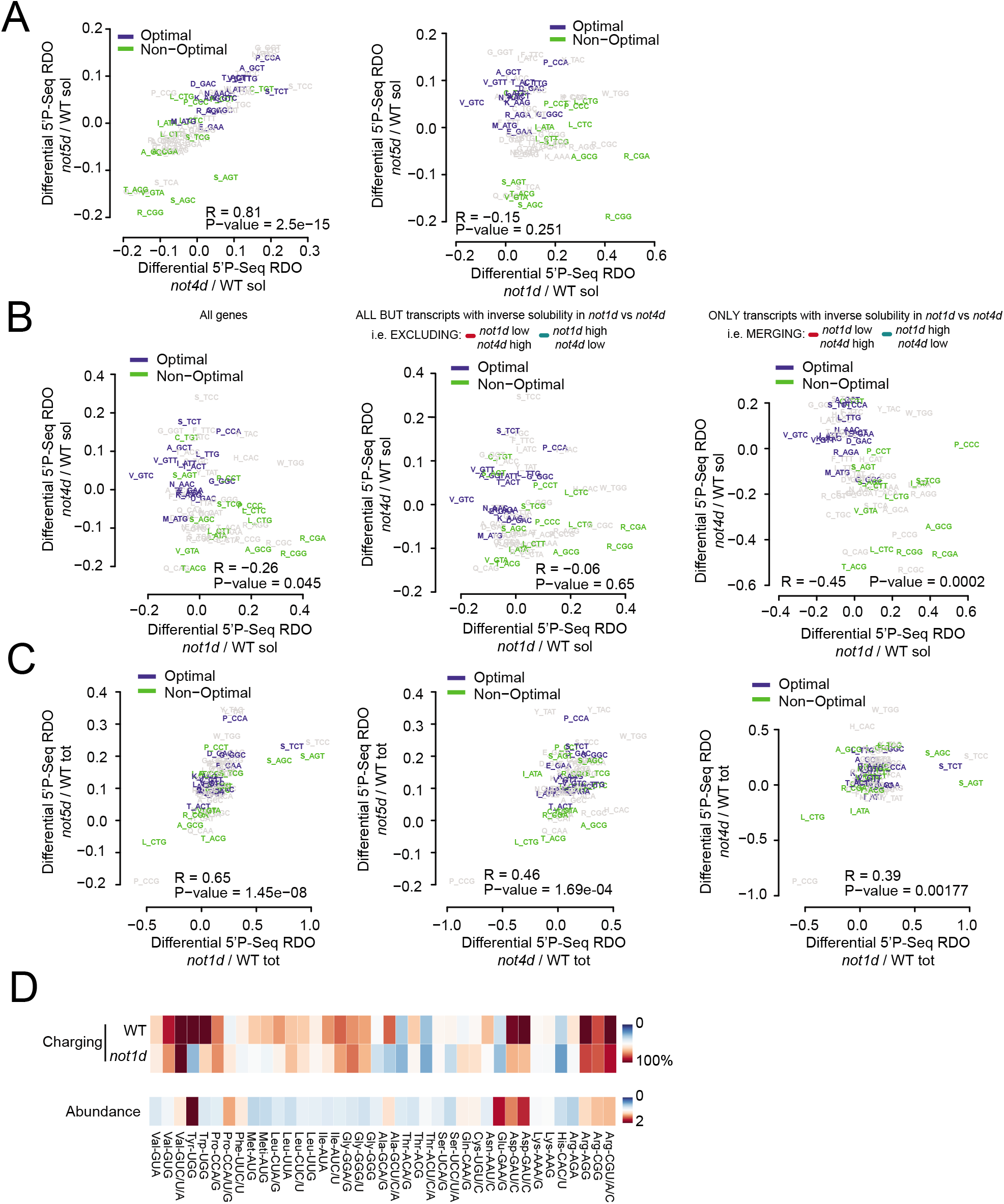
5’P-RDOs changes within the soluble RNA pool correlate with codon optimality upon Not4 and Not5 depletion. Scatterplot analyses comparing pairwise changes in 5’P-RDOs relative to WT in: **A**. soluble RNAs upon Not4 and Not5 (left) or Not1 and Not5 (right) depletion; **B**. soluble RNAs for Not1 and Not4 depletion for all mRNAs (left), all mRNAs excluding mRNAs of the red and green categories of **Figure 2C** (middle) or only for mRNAs of the red and green categories (right); **C**. for total RNAs upon Not1, Not4 and Not5 depletion. **D**. tRNA microarray analysis of aminoacyl-tRNA levels in Not1 depleted and corresponding wild-type strain (top lane) and the relative total tRNA abundance (bottom lane). The abundance was measured relative to the wild-type strain. The arrays are an average of three biological replicates. Confidence intervals between replicate 1 and 2, 1 and 3 and 2 and 3 were 97%, 98% and 98% for the charging arrays of the Not1 depleted cells, 97%, 97% and 97% for the abundance arrays of the Not1 depleted cells, 97%, 97% and 97% for charging arrays of the wild-type, and 95%, 94% and 98% for the abundance arrays, respectively. tRNA probes are depicted with their cognate codon and the corresponding amino acid.

For the total RNA pool, the 5’P-RDO changes observed immediately upon depletion of the Not proteins correlated, most importantly for Not1 and Not5 depletions and least importantly for Not1 and Not4 depletions, with some specific codons, e.g. Leu (CTG) and Pro (CCG) striking out (**Figure 5C**). 5’P-RDO changes in the total and soluble RNA pools after depletion of each respective Not protein did not show any significant correlation (**Figure S3B**).

5’P-RDOs increases at Ser codons were most dramatic upon Not1 depletion (**Figure 5C**). A change of metabolic flow from serine to alanine is known to occur during anoxia. Hence depletion of Not1 may have immediate changes on metabolism, which could be directly detected by the serine levels and tRNA charging, and would explain the dramatic 5’P-RDOs increases at Ser codons upon Not1 depletion. To test this, we determined the relative changes in the total RNA levels and the percentage of charged tRNA for each isoacceptor, before and following Not1 depletion. We observed a slight but insignificant decrease of the tRNA^Ser^ levels following Not1 depletion, but the levels of two seryl-tRNA^Ser^ isoacceptors markedly decreased (**Figure 5D**).

### mRNAs whose overexpression and solubility are inversely impacted by Not1 or Not4 depletion show relatively lower Not4 cross-linking

The inverse impacts of Not1 and Not4 on mRNA solubilities and 5’P-RDOs raises the question of the mechanism, and as to whether the regulation occurs via their mRNA binding. In previous work we defined Not1 mRNA binding by RNA immunoprecipitation (RIP) in wild type cells and in cells lacking Not5. We noted an interesting significant higher Not1 RIP in *not5Δ* for the mRNAs more soluble upon Not1 depletion (**Figure 6A**). This was not related to the overall increased size of the mRNAs bound by Not1 in the absence of Not5 (**Figure 2F**).

**Figure 6.**
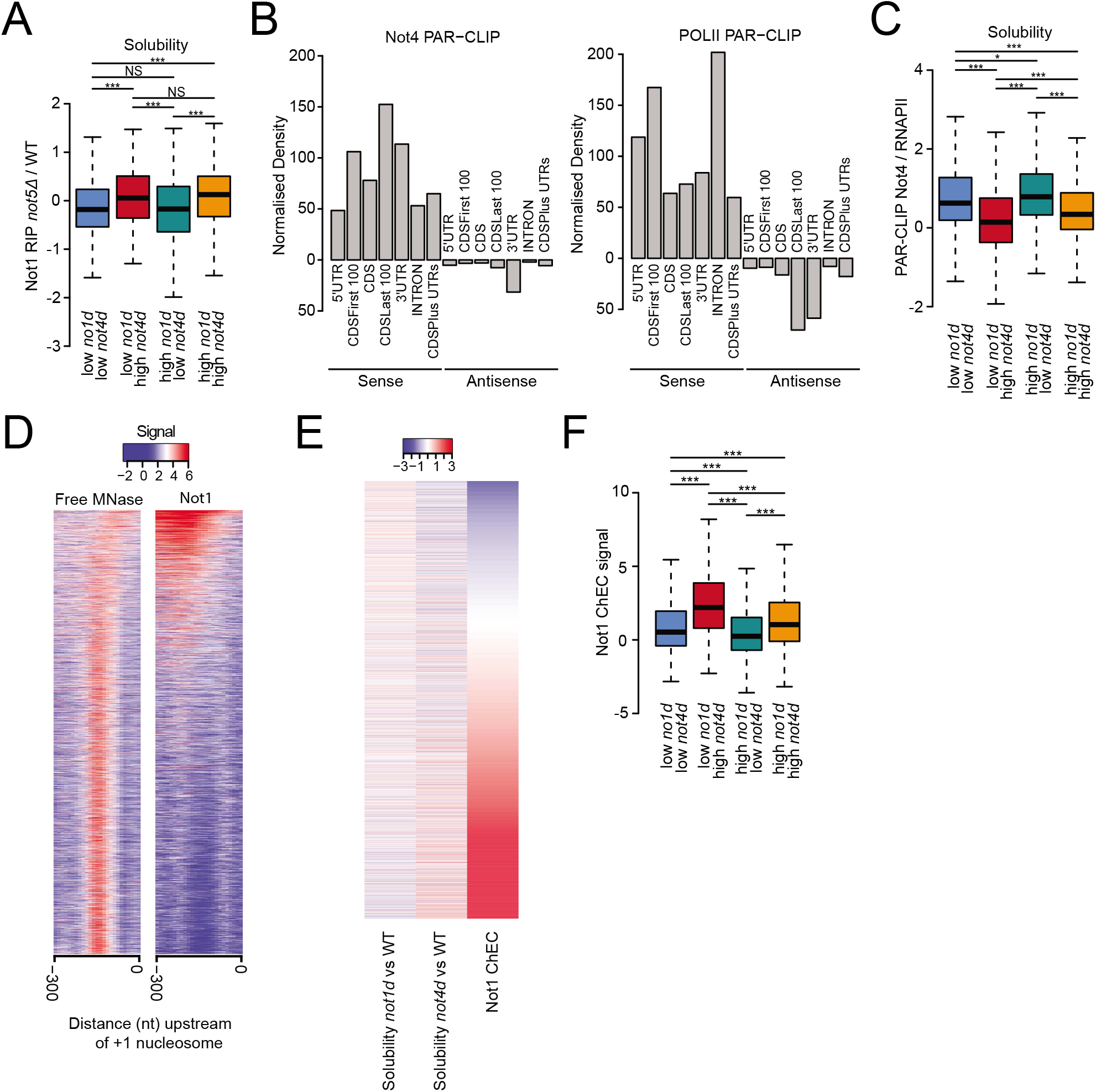
Not4 cross-linking correlates with changes in mRNA solubilities upon Not4 depletion and Not1 promoter association correlates with mRNAs less soluble upon Not1 depletion. **A**. Box plot comparing Not1 RIP in cells lacking Not5 for the categories of mRNAs color-coded in **Figure 2C**. Significance of differences is indicated at the top of the box plots - p-values are calculated using a two-sided Welch two sample t-test. **B**. Comparative cross-linking (normalized density) of Not4 (left) and RNAPII (right) to different mRNA sequences as indicated. **C**. Same as panel A, but comparing the relative Not4/RNAPII cross-linking to the different classes of mRNAs. **D**. Heat map comparing signal of free MNAse and Not1-MNase DNA cleavage events at promoter regions (ChEC). **E**. Heat map comparing Not1-ChEC signal and changes in mRNA solubilities upon Not1 and Not4 depletion. **F**. Box plot analysis of the Not1-ChEC signal in the different promoters of the genes encoding the different classes of mRNAs color-coded in **Figure 2C**. Significance of differences is indicated at the top of the box plots - p-values are calculated using a two-sided Welch two sample t-test.

We thus focused on Not4 whose association with RNAs has not yet been characterized. To define to which mRNAs Not4 binds *in vivo*, we used a photoactivatable ribonucleoside-enhanced crosslinking and immunoprecipitation (PAR-CLIP) approach (62). In two independent biological replicates we identified the positions of Not4 cross-linking to mRNAs, resulting in T to C transitions (**Table S3**). The replicates showed high correlation (**Figure S4A**). The reads were distributed throughout transcribed sequences, mapping to coding sequences (CDSs), 3’ and 5’ untranslated regions (UTRs), and introns, of the sense and anti-sense regions (**Figure 6B**, left). Overall reads correlated with T to C transitions on coding sequences as well as on introns (**Figure S4B**). There was a good correlation between cross-linking of Not4 and cross-linking of RNA polymerase II (RNAPII) (63) (**Figure 6B**, right) on all sense (**Figure S4C)**, and anti-sense (**Figure S4D**) sequences, suggesting Not4 binds mRNA ubiquitously during transcription. Despite the global correlation, we noticed some differences. For instance, RNAPII cross-linked more efficiently to introns and less to CDSs, whereas Not4 was cross-linked better to 3’ UTRs (**Figure 6B**), and the correlation was better for sense than for anti-sense regions (**Figure S4C** and **D**). Moreover, the patterns of Not4 and RNAPII cross-linking differed along mRNAs, with RNAPII cross-linking more prominent in 5’UTRs and at the very beginning of CDSs, but Not4 cross-linking instead more prominent on CDSs, most striking at the end of coding sequences and also more prominent in 3’UTRs (**Figure S4E**). Only few mRNAs (270) had at least 2-fold less cross-linking of Not4 than expected from the global correlation between Not4 and RNAPII cross-linking whereas 1222 mRNAs had more than 2-fold higher cross-linking of Not4 than expected (**Figure S4F**, in purple). GO-term analysis of this latter mRNA group revealed enriched categories, including “mitochondrion organization”, “response to oxidative stress”, “proteolysis involved in cellular protein catabolic processes” “protein complex biogenesis”, “protein folding”, “cytoplasmic translation” (**Figure S4G**), that have functionally been connected to Not4 in previous studies (17, 42, 49, 53, 64, 65).

We compared the mRNAs cross-linked to Not4 to the sets of mRNAs differentially solubilized described above (**Figure 2C**). mRNAs less soluble upon Not4 depletion showed higher Not4 cross-linking (**Figure 6C**, blue and green mRNAs), suggesting that Not4 mRNA binding plays a role for solubility of these mRNAs. Notably, this trend was also detectable, albeit to a lesser degree when Not4 cross-linking rather than relative Not4 to RNAPII cross-linking was considered (**Figure S4H**). Instead, it was not as significant for all comparisons when RNAPII (**Figure S4I**) or expression levels (RNA-Seq) (**Figure S4J**) were considered. This indicates that it is not only related to Not4 cross-linking to mRNAs according to expression levels, but to functions of Not4 after transcription.

### Not1 binds promoters broadly but with specificity related to mRNA solubility

Previous work has shown that Not5-dependent Not1 binding to ribosomal protein (RP) mRNAs occurs during transcription and regulates their translation in a manner dependent upon Not5 (48). Above we showed that the mRNAs more soluble upon Not1 depletion have higher Not1 RIP in *not5Δ*. We thus mapped Not1 promoter binding genome-wide to determine whether Not1 association with promoters correlated with the Not1-dependent fate of mRNAs in the cytoplasm, in particular their solubility. We fused Not1 to the micrococcal nuclease (MNase) at its own genomic locus, monitored chromatin cleavage events (ChEC) and compared them to the cleavage pattern by the free MNase. In total, 4923 promoters showed cleavages by Not1-MNase above the threshold of significance, and the pattern of Not1-MNase was specific compared to the pattern obtained with free MNase (**Table S4** and **Figure 6D**). Higher binding of Not1 at promoters correlated with lower solubility of mRNAs upon Not1 depletion and higher solubility of mRNAs upon Not4 depletion (**Figure 6E**). In particular, the promoters driving transcription of mRNAs showing lower solubility upon Not1 depletion but higher following Not4 depletion (red mRNAs), showed higher Not1 promoter binding (**Figure 6F**). These results suggest that the cytoplasmic fates of mRNAs defined by Not1 and Not4 in opposing manner are likely set in the nucleus during transcription by Not1 promoter binding and resulting lower Not4 mRNA cross-linking.

## Discussion

### Solubility as a mechanism by which Not proteins regulate co-translation dynamics

Several recent studies provide evidence that the Not proteins are key for codon-optimality related changes in mRNA stability (50, 66). A beautiful structure of Not5 associated with the translating ribosome corroborates the idea that the Ccr4-Not complex can monitor codon optimality, but a mechanism linking this ribosome docking to control of mRNA decay is elusive. Our work has indicated that ribosome dwelling occupancy evaluated by ribosome profiling (i.e. valid for the soluble mRNA pool) is regulated according to codon-optimality by the Not subunits of the Ccr4-Not complex. We proposed that this could occur via the dynamic formation and dissolution of Not condensates during translation (17). In this current study we tested the model further by comparing soluble mRNAs to the total mRNA pool that includes all insoluble mRNAs. We find that soluble mRNAs differ from the insoluble mRNAs by higher relative degradation and lower content in ribosome dwelling at non-optimal codons, and that Not1 and Not4, both acting with Not5, inversely modulate solubility for different mRNAs These findings indicate that modulation of solubility is the mechanism by which Not5-ribosome binding can regulate mRNA turnover and translation dynamics according to codon optimality.

### Soluble and insoluble RNAs pools have distinguishable co-translational dynamics

We show that solubility of mRNAs is a determining factor in regulating co-translation dynamics. Indeed, soluble mRNAs show more relative degradation but less detectable co-translational decay compared to insoluble mRNAs, namely all mRNAs that are not soluble without distinction of the different types of insoluble mRNAs. In soluble fractions ribosome movement is faster than the action of the 5’-3’ exonuclease activity. Alternatively, or in addition, mechanisms other than co-translational decay might generate decay intermediates. These could be No-Go-Decay (NGD) that can generate 5’ to 3’ decay intermediates upstream of collided ribosomes (67), and more importantly post-translational decay that is not marked by ribosome dwelling.

Soluble and insoluble mRNAs also exhibit a different distribution of 5’P decay intermediates along CDSs most likely due to differences in ribosome dwelling and hence velocity. Indeed, we show that ribosomes dwelling at non-optimal codons are more prominent in the insoluble RNA pool. Alternatively, the delay of 5’ to 3’ decay initiation after the last trailing ribosome, indicated by higher 5’P-RDOs at non-optimal codons for the second half of CDSs compared to the first half, is less significant for soluble mRNAs, since RNA-Seq reads relative to 5’P-Seq reads in the first half compared to the second half of CDSs are higher.

### mRNA solubility is inversely regulated by Not1 and Not4, both working with Not5

In previous work we proposed that tethering of mRNAs by Not proteins to condensates, thus to insoluble RNA pools, in particular at start and at non-optimal codons, modulates translation elongation dynamics (17). In line with this model, and based on the results discussed here, depletions of the Not proteins increase overall the relative degradation and, in addition, three-nucleotide periodicity overall. This implies a more prominent role of the Not proteins for general decay than for co-translational decay. It is consistent with Not proteins being subunits of the major eukaryotic deadenylase complex (22) and with the role of Not4 for a bypass quality control pathway (68).

While these findings indicate that the Not proteins act together and in a similar manner, several observations contradict this simple interpretation. First, depletion of Not4 or Not5 results in higher, but that of Not1 lower, relative degradation for soluble mRNAs. Higher levels of 5’P decay intermediates in the second CDS half are observed most significantly upon depletion of Not1 for soluble mRNAs, but upon depletion of Not4 for total mRNAs. Importantly, mRNA solubility was mildly altered by depletion of Not5, but inversely impacted by the depletions of Not1 and Not4 (see model on **Figure 7**). In particular, mRNAs more soluble upon Not4 depletion (red category) have a relatively lower content in non-optimal codons and the 5’P-RDO changes for soluble mRNAs that correlate upon Not4 and Not5 depletion and correlate with codon optimality, can be related to the mRNAs with low non-optimal codons becoming soluble. Instead, they become insoluble upon Not1 depletion, hence 5’P-RDO changes tend to be inverse.

**Figure 7.**
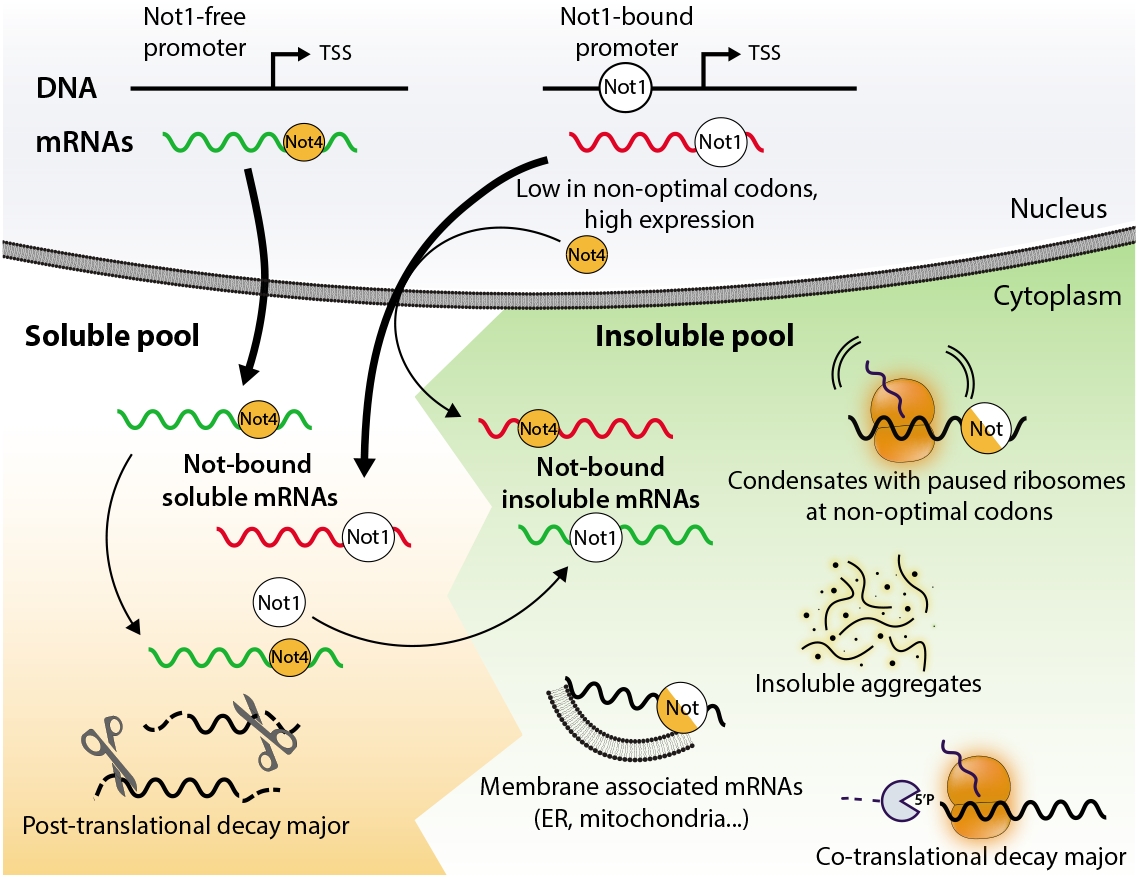
Model for the opposing regulation of mRNA solubility by Not1 and Not4. mRNAs exhibit diverse solubilities, and mRNAs that are not soluble can be either in functional condensates, associated with membranes or in cytosolic aggregates. Not1 and Not4 have opposing effects on mRNA solubility and solubility of some mRNAs (designated red) is promoted by Not1, whereas for another set (green), solubility is promoted by Not4. mRNAs that are less soluble upon Not1 depletion but more upon Not4 depletion (red), are transcribed from genes with high Not1 recruited at their promoters. Not4, also co-transcriptionally recruited to mRNAs, opposes their solubilization. These mRNAs have low non-optimal codon content and are highly expressed. They show high Not1 binding in *not5Δ* (RIP) background and Not4 cross-linking is lower compared to the mRNAs that instead are less soluble upon Not4 depletion but more upon Not1 depletion (green mRNAs). These latter mRNAs are generally more soluble and they are less likely to have co-transcriptional recruitment of Not1. They remain soluble in a Not4-dependent manner. Solubility of these mRNAs is compromised by the depletion of Not4, possibly because Not1 recruitment is thereby enabled, since depletion of Not1 instead increases their solubility. Note that Not1 and Not4 are placed on mRNAs to indicate functional interaction, but does not to infer direct mRNA binding.

In contrast to Not1 and Not4 depletion, Not5 depletion showed only minor effects on mRNA solubility. We expect that Not5 association with ribosomes is key for regulations by Not1 and Not4 since recent work features a key role of Not5 in monitoring codon optimality via its binding to post-translocation ribosomes (50). Indeed, Not5 works together with Not1 for mRNAs solubilized upon Not1 depletion (green category) and 5’P-RDO changes for total RNAs correlate best upon Not1 and Not5 depletion, but, as mentioned above, with Not4 for mRNAs solubilized upon Not4 depletion (red category). Thus, Not5 works with both Not1 and Not4 that have opposing roles, and these opposite effects most likely cancel each other upon Not5 depletion.

### Targets of Not1 regulation are conserved

A recent study has investigated the global effects of CNOT1 knockdown in human cell lines (69). An interesting parallel can be seen between the classes of mRNAs regulated in human by CNOT1, either at the mRNA stability level of at the translation efficiency level, and those regulated by Not1 in yeast. Indeed, in yeast shorter mRNAs are less soluble upon Not1 depletion. Such mRNAs would be expected to be more stable in Not1 knockdown and this correlates with the finding that in CNOT1 knockdown half-life of shorter mRNAs increases. In yeast, mRNAs less soluble upon Not1 knockdown are enriched for mRNAs translated at the ER. In human cells, upon CNOT1 depletion, ER-targeted mRNAs were enriched within mRNAs with reduced translation efficiency, calculated as a ratio of ribosome footprints determined by ribosome profiling relative to total RNA. Ribosome footprinting can only evaluate soluble RNA fractions, thus if these mRNAs are less soluble upon CNOT1 depletion, one would expect a drop in translation efficiency. In an opposite manner, in yeast, Not1 depletion increases solubility of mitochondrial mRNAs and CNOT1 knockdown increased their translation efficiency in human cells, a phenotype that would be observed if the mRNAs are more soluble. These comparisons indicate that inherent mRNA characteristics of some gene groups regulated by Not1 are conserved from yeast to human.

### Regulation of mRNA solubility is set during transcription

It might appear counter-intuitive that two subunits of the same complex have opposing roles. However, having two opposing factors working in the context of a single multi-subunit complex might be essential to fine-tune co-translational dynamics. It should also be noted that in human cells CNOT4 is not a stable subunit of the Ccr4-Not complex. It raises the question as to how the opposing roles of Not1 and Not4 are set. The inverse regulation by Not1 and Not4 is importantly correlated with levels of Not1 promoter binding and Not4 cross-linking, and our data clearly indicates that Not4 association with mRNAs occurs during transcription. Hence, mRNA solubility appears to be set already during transcription. It could be that high Not1 promoter binding can result in higher Not1 association with newly produced mRNAs and thus lower Not4 cross-linking, whereas if less Not1 is present at the promoter, higher Not4 association with newly produced mRNAs can occur (see model on **Figure 7**). After translation onset, the roles of Not1 and Not4 may dynamically interchange during translation elongation, in line with our previous finding that Not condensates are dynamic (17). Not condensates may be more or less insoluble according to their complexity or their association for instance with membranes.

Not1 appears to be important for solubility of highly expressed mRNAs and important for cytoplasmic translation, not only of ribosomal protein mRNAs, whose solubilities are inversely regulated by Not4, but also of mRNAs encoding tRNA processing and rRNA modification enzymes that are important for the production of a functional translation machinery. It could be that Not1 plays a role to prevent aggregation or condensation of such mRNAs during translation and/or to release such mRNAs from condensates. Indeed, a role of Not1 to solubilize Dhh1 condensates has been demonstrated (70). Instead, mRNAs with higher Not4 cross-linking and lower Not1 at promoters tend to be more soluble (blue and green categories) suggesting that higher association of Not4 with mRNAs counteracts a more general effect of Not1 to contribute to mRNA insolubility. Depletion of Not1 and Not4 also have similar effects, rendering shorter mRNAs less soluble but solubilizing longer mRNAs. Length of mRNAs may counterbalance the effect of Not proteins co-transcriptionally recruited to mRNAs by co-translational recruitment of additional factors.

## Acknowledgments

Not applicable

## Declarations

### Ethics approval and consent to participate

Not applicable

### Consent for publication

Not applicable

### Availability of data and materials

tRNA microarray data are accessible under the accession number GSE190658 (the following token: opgbkqmujtylnwz, is for the reviewers, if they want to have a look at the primary data) in the Gene Expression Omnibus (GEO) database, 5’P-Seq data under GSE193912. All code used in the manuscript is available https://github.com/georgeallenunige/NotTranslation

### Author contributions

GEA performed 5’P-Seq and RNA-Seq analyses, BW prepared the RNAs. OOP prepared strains for Par-CLIP. ZV did the Not1-RIP and the ChEC-Seq with BA and DD in the laboratory of DS. MZ prepared the degron strains. SH made the 5’P libraries and initial analyses with VP, CP performed and analyzed the tRNA microarrays and analysis with ZI. PS, SEL and JC did the Not4 PAR-CLiP. BW prepared figures and contributed to manuscript design. ZI, VP and MC analyzed data, supervised the work and wrote the manuscript.

### Authors’ information

Not applicable

## Materials and Methods

### Strains, plasmids and culture conditions

The Not4 and Not5 degron strains (13771 and 13772) were created in strain MY13472 (YDR10, kind gift from from David Shore) and PCR amplification of a 9Myc-NATMX4 cassette with Not4- and Not5- specific primers using plasmid pE641. The Not1 degron strain (13517) has already been described (17). The strains were verified by PCR. Not protein depletions were obtained by addition of auxin (3-indoleacetic acid, Sigma-Aldrich I2886, stock solution at 250 mM in EtOH) at 1 mM final for 15 min to exponentially growing cells diluted to OD_600_ 0.3 after an overnight culture in glucose rich medium (YPD), when they reached OD_600_ 0.8. Equivalent amounts of EtOH 100% were added for the control. All experiments were performed with cells growing in YPD. The strain expressing the Not4-HTB fusion (MY11050) was generated with Not4-specific primers by PCR using pE557, the strains expressing the Not1-MNase fusion (MY12601) was generated with Not1-specific primers by PCR using pE611 respectively. The free MNase strain (MY13783) was YMC08 (kind gift from David Shore).

### RNA preparation

Total RNA was prepared either by the hot acid phenol method (71) or cells were prepared and lysed as for polysome profiling (42) and total RNA was prepared from the lysate. Briefly, for total RNA, cell pellets from 30 ml of exponentially growing yeast were resuspended with 400 μl of TES buffer (10 mM Tris HCl pH 7.5, 10 mM EDTA and 0.5 % SDS) to which 400 μl of acid phenol were added. After vortexing and incubation for 10 min at 65°C, RNA was extracted. For soluble RNA, cell pellets from 30 ml of exponentially growing yeast were resuspended in 400 μl of lysis buffer (20 mM Hepes 20 mM KCl, 10 mM MgCl2 1% triton, 1mM PMSF, 1 mM DTT supplemented with a cocktail of protease inhibitors and with 0.1 mg/ml of cycloheximide) to which 200 μl of glass beads were added. Cells were vortexed at 4°C for 15 min and spun at 15000 K for 1 min. The supernatant was transferred to a new tube and spun a further 20 min at 4°C. The supernatant was combined with 400 μl of acid phenol for further RNA extraction as for the total RNA pool.

### Protein extraction and analysis

Total protein was prepared by post-alkaline lysis and analyzed by western blotting with antibodies to Not1, Not4 and Not5 that were our own polyclonal antibodies previously described (72, 73). Antibodies to Myc were commercial (Sigma M5546).

### 5’P-Seq

RNA was prepared from 50 ml of exponentially growing cells in YPD and treated or not with auxin. HT-5PSeq libraries were generated as reported (74) with minor modifications. In brief, 15μg total RNA, containing 5% total RNA from Schizosaccharomyces pombe as spike-in, was used. Each sample was spited in two. One part was used for preparing conventional HT-5PSeq libraries and the other part for was random fragmented prior to the preparation of HT-5PSeq libraries (negative control).

For HT-5PSeq Libraries: 7.5 μg RNA was ligated over night at 16°C to r5P_RNA_MPX oligo (CrArCrGrArCrGrCrUrCrUrUrCrCrGrArUrCrU rXrXrXrXrXrX rNrNrNrNrNrNrNrN) carrying a sample barcode (rX) and unique molecular identifiers (rN). Ligase was deactivated using 5mM EDTA and heat at 65°C for 10 minutes (up to X individual barcoded RNA ligations were pooled) and subsequent purified using 1.8x volumes of RNAClean XP beads (Beckman Coulter). Ligated RNA was then reverse transcribed using random hexamer (5Pseq-RT, GTGACTGGAGTTCAGACGTGTGCTCTTCCGATCTNNNNNN, 20 μM) and oligo-dT (5Pseq-dT, GTGACTGGAGTTCAGACGTGTGCTCTTCCGATCTTTTTTTTTTT at 0.05 μM) oligos to prime. After, remaining RNA was degraded using NaOH. Ribosomal RNA was removed using previously described rRNA DNA oligo depletion mixes, following a duplex-specific nuclease (DSN, Evrogen) digestion. rRNA depleted cDNA was amplified by PCR (17 cycles) and final product was enriched for fragments with the range of 300-500 nt using Ampure XP.

Size selected HT-5P Libraries were quantified by fluorescence (Qubit, Thermo Fisher), size estimated using an Agilent Bioanalyzer and sequenced using a NextSeq500 Illumina sequencer (75 cycles High output kit).

### Not4 PAR-CLIP

Cells expressing Not4 tagged at the C-terminus with the HTB tag (75) from its endogenous locus and endogenous promoter were grown in duplicates in the presence of 4-thiouracil and then UV-irradiated at 365 nm to cross-link proteins with RNA. Not4 was purified under denaturing conditions and libraries were prepared from the co-purified RNA and sent for deep sequencing as described (63).

### Chromatin endogenous cleavage (ChEC-Seq)

Not1 ChEC-Seq experiments was essentially performed as previously described (76) (Zentner et al. 2015) with the following modifications. Cells in which MNase was fused at the C-terminus of the endogenous *NOT1* or *NOT5* genes were used to determine Not1 and Not5 binding. Cells in which MNase was placed under the control of *REB1* promoter were used as a control. One sample corresponds to 12 ml of culture at OD_600_ = 0.7. Cells were washed twice with buffer A (15 mM Tris 7.5, 80 mM KCl, 0.1 mM EGTA, 0.2 mM spermine, 0.5 mM spermidine, 1xRoche EDTA-free mini protease inhibitors, 1 mM PMSF) and resuspended in 200 μl of buffer A with 0.1% digitonin. The cells were incubated for 5 min at 30°C. Then, MNase action was induced by addition of 5 mM CaCl_2_ and stopped at 150 seconds for Not1 and Not5 ChEC-seq and 20 minutes for Mnase under the control of REB1 promoter by adding EGTA to a final concentration of 50 mM. DNA was purified using MasterPure Yeast DNA purification Kit (Epicentre) according to the manufacturer’s instruction. Large DNA fragments were removed by a 5-min incubation with 2.5x volume of AMPure beads (Agencourt) after which the supernatant was kept, and MNase-digested DNA was precipitated using isopropanol. Libraries were prepared using NEBNext kit (New England Biolabs) according to the manufacturer’s instructions. Before the PCR amplification of the libraries small DNA fragments were selected by a 5-minute incubation with 0.9x volume of the AMPure beads after which the supernatant was kept and incubated with the same volume of beads as before for another 5 min. After washing the beads with 80% ethanol the DNA was eluted with 0.1x TE and PCR was performed. Adaptor dimers were removed by a 5-min incubation with 0.8x volume of the AMPure beads after which the supernatant was kept and incubated with 0.3x volume of the beads. The beads were then washed twice with 80% ethanol and DNA was eluted using 0.1x TE. The quality of the libraries was verified by running an aliquot on a 2% agarose gel. Libraries were sequenced using a HiSeq 2500 machine in single-end mode. To analyze the Not1- and Not5-MNase binding pattern, read ends were considered to be MNase cuts and were mapped to the genome (sacCer3 assembly) using HTSstation (David et al. 2014).

### tRNA microarrays

To determine the fraction of aminoacyl-tRNAs we followed the procedure described in (77). For this, total RNA was isolated in mild acidic conditions (pH 4.5) which preserves the aminoacyl-moiety. Each sample was split into two aliquots and one was oxidized with periodate to which the changed tRNAs remain intact and following subsequent deacylation (100 mM Tris (pH 9.0) at 37°C for 45 min) was hybridized to Cy3-labeled RNA/DNA stem-loop oligonucleotide. The second aliquot was deacylated to receive the total tRNA and hybridized to Atto647-labeled RNA/DNA stem-loop oligonucleotide. Both aliquots were analyzed on the same tRNA microarrays and the ratio of the Cy3 to Atto647 signal provides the fraction of aminoacyl-tRNA for each isoacceptor.

For tRNA abundance, total RNA was isolated at alkaline pH to simultaneously deacylate all tRNAs. tRNAs isolated from Not1 depleted cells were labeled with Cy3-labeled RNA/DNA stem-loop oligonucleotide and were hybridized on the same microarray with tRNAs isolated from the wild-type strain and labeled with Att647-labeled RNA/DNA stem-loop oligonucleotide. The arrays were normalized to spike-in standards, processed and quantified with in-house python scripts.

### Bioinformatic analyses

#### 5’P-Seq and RNA-Seq

Sequencing files were demultiplexed using bcl2fastq v2.20.0.422 (one mismatch, minimum length 35 nt), and adapters were trimmed using cutadapt 2.3. (78) at default settings, allowing one mismatch and minimum read length of 35nt. In addition to standard illumine dual index (i5, i7), the inline sample and UMI barcode was analyzed using Umitools. Reads were mapped to the concatenated genome of *S. cerevisiae* (R64-1-1) and *S. pombe* (ASM294v2) using STAR.

Second read enables to splits reads between oligo-dT or random primer. That information was not used in the current analysis. CDS positions were defined with Ensembl gff version 94 for of *S. cerevisiae* (R64-1-1). Counts in *S. cerevisiae* were calculated by aggregating RNA-Seq reads and 5’P-Seq 5’-ends, overlapping CDS positions. Differential expression was performed using DESeq2 (79).

#### Solubility

We define this as the log fold change produced by DESeq2, dividing RNA-Seq counts for the soluble fraction in a given sample by the corresponding counts for the total fraction of the same sample.

#### Relative degradation

Spike-in *S.pombe* data was used in the calculation of relative degradation. We define it as the log fold change produced by DESeq2, comparing counts in 5’P-Seq to their corresponding RNA-Seq sample using estimateSizeFactors on counts mapping to *S. pombe* to adjust for spike-in. Enrichment is calculated using a hypergeometric test for over-representation of hits in defined gene set for GO SLIM categories for *S. cerevisiae*.

To calculate 5’P-Seq pausing scores, equivalent to A site ribosome dwelling occupancy, the mean depth was calculated 17nt upstream of each codon type for each strain in the regions or transcripts of interest. In all cases, the values are normalised to the mean depth over all codons for the regions or transcripts included in the calculation. Where two conditions are compared, the differential RDO is calculated as the log2 fold change of these normalised values for each codon.

For 5’P-Seq, metagenes at start and stop are calculated by aggregating the depth of 5’ ends at each position relative to start or stop for every CDS and normalising each by the total depth per million genome-wide. These values are shifted 17nt downstream for equivalency with the A-site position, in the case of co-translational decay.

For scaled metagenes, every CDS was split into 100 equal bins and the mean depth of 5’ ends of RNA-Seq and 5’P-Seq was calculated for each bin. This was averaged over all transcripts of interest and normalised to the mean depth over all nucleotides in this transcript group.

#### PAR-CLIP analyses

FASTQ files were adapter stripped, using cutadapt (parameters: -a AGATCGGAAGAGCACACGTCTGAACTCCAGTC --minimum-length=13 --quality-cutoff=2) and then mapped using bowtie (80) to sacCer3 (parameters: -v 2 -m 10 --best –strata). High confidence T to C transitions in the cDNA sequence defining the sites of cross-linked 4-thiouracil residues were identified using wavClusteR (81). These were then normalised to the rate of T bases for the regions of interest to give a normalised density value for cross-linking.

#### ChEC-seq analyses

FASTQ files were adapter stripped, using cutadapt (parameters: -a GATCGGAAGAGCACACGTCTGAACTCCAGTCA --minimum-length=20 --quality-cutoff=2) and then mapped using bowtie2 (82) to sacCer3 (parameters: -v 2 -m 10 --best – strata). Positions of the +1 nucleosome associated with each gene were taken from the Saccharomyces Genome Database and read counts overlapping the promoter binding region 400bp upstream and 100bp downstream of these position were calculated and normalized to RPKMs. To find the Not1 ChEC signal, the log2 fold change (LFC) of the promoter binding region RPKMs were taken over free MNase. These values were mode-centred to zero (the mode of the LFCs was estimated by fitting a log-normal distribution using ‘fitdistr’ from the R package MASS).

#### Ribo-Seq

Values for Ribo-Seq RPKMs were calculated as in our previous paper [17].

### Statistical tests

Reported correlations were the Pearson’s product-moment correlation coefficient and were used as test statistics to generate the associated p-value by t-test. All correlations and correlation tests were performed on groups of at least 30 in size. Enrichment of gene sets is defined via FDR after Benjamini-Hochberg adjustment from p-values generated using a hypergeometric test. All t-tests were performed on sample sizes of 30 or higher, where the central limit theorem applies regarding the normality assumption. We use Welch’s *t*-test in all cases, rather than the Student’s t-test, resulting in more conservative *p*-value, which is more reliable where variances and sample sizes are unequal. In one case, we used a Wilcoxon rank sum test (83) to compare two samples as a non-parametric proxy for a *t*-test, so as to avoid any possible breach of the normality distribution assumption, since both samples had a size of 15 (comparing WT RDOs Total/Soluble for optimal and non-optimal groups).

## References

1. Corbett AH. Post-transcriptional regulation of gene expression and human disease. Curr Opin Cell Biol. 2018;52:96–104.

2. Sunnerhagen P. Cytoplasmatic post-transcriptional regulation and intracellular signalling. Mol Genet Genomics. 2007;277(4):341–55.

3. Filipowicz W, Bhattacharyya SN, Sonenberg N. Mechanisms of post-transcriptional regulation by microRNAs: are the answers in sight? Nat Rev Genet. 2008;9(2):102–14.

4. Schaefke B, Sun W, Li YS, Fang L, Chen W. The evolution of posttranscriptional regulation. Wiley Interdiscip Rev RNA. 2018:e1485.

5. Bresson S, Shchepachev V, Spanos C, Turowski TW, Rappsilber J, Tollervey D. Stress-Induced Translation Inhibition through Rapid Displacement of Scanning Initiation Factors. Mol Cell. 2020;80(3):470–84 e8.

6. Advani VM, Ivanov P. Translational Control under Stress: Reshaping the Translatome. Bioessays. 2019;41(5):e1900009.

7. Sonenberg N, Hinnebusch AG. Regulation of translation initiation in eukaryotes: mechanisms and biological targets. Cell. 2009;136(4):731–45.

8. Richter JD, Coller J. Pausing on Polyribosomes: Make Way for Elongation in Translational Control. Cell. 2015;163(2):292–300.

9. Riba A, Di Nanni N, Mittal N, Arhne E, Schmidt A, Zavolan M. Protein synthesis rates and ribosome occupancies reveal determinants of translation elongation rates. Proc Natl Acad Sci U S A. 2019;116(30):15023–32.

10. Knight JRP, Garland G, Poyry T, Mead E, Vlahov N, Sfakianos A, et al. Control of translation elongation in health and disease. Dis Model Mech. 2020;13(3).

11. Stahl T, Hummer S, Ehrenfeuchter N, Mittal N, Fucile G, Spang A. Asymmetric distribution of glucose transporter mRNA provides a growth advantage in yeast. EMBO J. 2019;38(10).

12. Ma W, Mayr C. A Membraneless Organelle Associated with the Endoplasmic Reticulum Enables 3’UTR-Mediated Protein-Protein Interactions. Cell. 2018;175(6):1492–506 e19.

13. Das S, Vera M, Gandin V, Singer RH, Tutucci E. Intracellular mRNA transport and localized translation. Nat Rev Mol Cell Biol. 2021;22(7):483–504.

14. Cioni JM, Lin JQ, Holtermann AV, Koppers M, Jakobs MAH, Azizi A, et al. Late Endosomes Act as mRNA Translation Platforms and Sustain Mitochondria in Axons. Cell. 2019;176(1–2):56–72 e15.

15. Decker CJ, Parker R. P-bodies and stress granules: possible roles in the control of translation and mRNA degradation. Cold Spring Harb Perspect Biol. 2012;4(9):a012286.

16. Mateju D, Eichenberger B, Voigt F, Eglinger J, Roth G, Chao JA. Single-Molecule Imaging Reveals Translation of mRNAs Localized to Stress Granules. Cell. 2020;183(7):1801–12 e13.

17. Allen GE, Panasenko OO, Villanyi Z, Zagatti M, Weiss B, Pagliazzo L, et al. Not4 and Not5 modulate translation elongation by Rps7A ubiquitination, Rli1 moonlighting, and condensates that exclude eIF5A. Cell Rep. 2021;36(9):109633.

18. Panasenko OO, Somasekharan SP, Villanyi Z, Zagatti M, Bezrukov F, Rashpa R, et al. Co-translational assembly of proteasome subunits in NOT1-containing assemblysomes. Nat Struct Mol Biol. 2019;26(2):110–20.

19. Lee CD, Tu BP. Glucose-Regulated Phosphorylation of the PUF Protein Puf3 Regulates the Translational Fate of Its Bound mRNAs and Association with RNA Granules. Cell Rep. 2015;11(10):1638–50.

20. Ingolia NT, Ghaemmaghami S, Newman JR, Weissman JS. Genome-wide analysis in vivo of translation with nucleotide resolution using ribosome profiling. Science. 2009;324(5924):218–23.

21. Pelechano V, Wei W, Steinmetz LM. Widespread Co-translational RNA Decay Reveals Ribosome Dynamics. Cell. 2015;161(6):1400–12.

22. Collart MA. The Ccr4-Not complex is a key regulator of eukaryotic gene expression. Wiley Interdiscip Rev RNA. 2016.

23. Collart MA, Struhl K. CDC39, an essential nuclear protein that negatively regulates transcription and differentially affects the constitutive and inducible HIS3 promoters. Embo J. 1993;12(1):177–86.

24. Collart MA, Struhl K. NOT1(CDC39), NOT2(CDC36), NOT3, and NOT4 encode a global-negative regulator of transcription that differentially affects TATA-element utilization. Genes Dev. 1994;8(5):525–37.

25. Denis CL, Malvar T. The *CCR4* gene from *Saccharomyces cerevisiae* is required for both nonfermentative and *spt*-mediated gene expression. Genetics. 1990;124:283–91.

26. Denis CL, Chiang YC, Cui Y, Chen J. Genetic evidence supports a role for the yeast CCR4-NOT complex in transcriptional elongation. Genetics. 2001;158(2):627–34.

27. Deluen C, James N, Maillet L, Molinete M, Theiler G, Lemaire M, et al. The Ccr4-not complex and yTAF1 (yTaf(II)130p/yTaf(II)145p) show physical and functional interactions. Mol Cell Biol. 2002;22(19):6735–49.

28. Kruk JA, Dutta A, Fu J, Gilmour DS, Reese JC. The multifunctional Ccr4-Not complex directly promotes transcription elongation. Genes & development. 2011;25(6):581–93.

29. Dutta A, Babbarwal V, Fu J, Brunke-Reese D, Libert DM, Willis J, et al. Ccr4-Not and TFIIS Function Cooperatively To Rescue Arrested RNA Polymerase II. Mol Cell Biol. 2015;35(11):1915–25.

30. Babbarwal V, Fu J, Reese JC. The Rpb4/7 module of RNA polymerase II is required for carbon catabolite repressor protein 4-negative on TATA (Ccr4-not) complex to promote elongation. J Biol Chem. 2014;289(48):33125–30.

31. Tucker M, Staples RR, Valencia-Sanchez MA, Muhlrad D, Parker R. Ccr4p is the catalytic subunit of a Ccr4p/Pop2p/Notp mRNA deadenylase complex in *Saccharomyces cerevisiae*. EMBO J. 2002;21:1427–36.

32. Tucker M, Valencia-Sanchez MA, Staples RR, Chen J, Denis CL, Parker R. The transcription factor associated Ccr4 and Caf1 proteins are components of the major cytoplasmic mRNA deadenylase in Saccharomyces cerevisiae. Cell. 2001;104(3):377–86.

33. Doidge R, Mittal S, Aslam A, Winkler GS. Deadenylation of cytoplasmic mRNA by the mammalian Ccr4-Not complex. Biochem Soc Trans. 2012;40(4):896–901.

34. Wahle E, Winkler GS. RNA decay machines: Deadenylation by the Ccr4-Not and Pan2-Pan3 complexes. Biochimica et biophysica acta. 2013.

35. Wilczynska A, Gillen SL, Schmidt T, Meijer HA, Jukes-Jones R, Langlais C, et al. eIF4A2 drives repression of translation at initiation by Ccr4-Not through purine-rich motifs in the 5’UTR. Genome Biol. 2019;20(1):262.

36. Eulalio A, Huntzinger E, Nishihara T, Rehwinkel J, Fauser M, Izaurralde E. Deadenylation is a widespread effect of miRNA regulation. Rna. 2009;15(1):21–32.

37. Chen CY, Zheng D, Xia Z, Shyu AB. Ago-TNRC6 triggers microRNA-mediated decay by promoting two deadenylation steps. Nat Struct Mol Biol. 2009;16(11):1160–6.

38. Goldstrohm AC, Seay DJ, Hook BA, Wickens M. PUF protein-mediated deadenylation is catalyzed by Ccr4p. J Biol Chem. 2007;282(1):109–14.

39. Enwerem, III, Elrod ND, Chang CT, Lin A, Ji P, Bohn JA, et al. Human Pumilio proteins directly bind the CCR4-NOT deadenylase complex to regulate the transcriptome. RNA. 2021;27(4):445–64.

40. Mishima Y, Fukao A, Kishimoto T, Sakamoto H, Fujiwara T, Inoue K. Translational inhibition by deadenylation-independent mechanisms is central to microRNA-mediated silencing in zebrafish. Proc Natl Acad Sci U S A. 2012;109(4):1104–9.

41. Alhusaini N, Coller J. The deadenylase components Not2p, Not3p, and Not5p promote mRNA decapping. RNA. 2016:709–21.

42. Panasenko OO, Collart MA. Presence of Not5 and ubiquitinated Rps7A in polysome fractions depends upon the Not4 E3 ligase. Molecular microbiology. 2012;83(3):640–53.

43. Panasenko O, Landrieux E, Feuermann M, Finka A, Paquet N, Collart MA. The yeast Ccr4-Not complex controls ubiquitination of the nascent-associated polypeptide (NAC-EGD) complex. J Biol Chem. 2006;281(42):31389–98.

44. Takehara Y, Yashiroda H, Matsuo Y, Zhao X, Kamigaki A, Matsuzaki T, et al. The ubiquitination-deubiquitination cycle on the ribosomal protein eS7A is crucial for efficient translation. iScience. 2021;24(3):102145.

45. Matsuki Y, Matsuo Y, Nakano Y, Iwasaki S, Yoko H, Udagawa T, et al. Ribosomal protein S7 ubiquitination during ER stress in yeast is associated with selective mRNA translation and stress outcome. Sci Rep. 2020;10(1):19669.

46. Dimitrova LN, Kuroha K, Tatematsu T, Inada T. Nascent peptide-dependent translation arrest leads to Not4p-mediated protein degradation by the proteasome. J Biol Chem. 2009;284(16):10343–52.

47. Villanyi Z, Ribaud V, Kassem S, Panasenko OO, Pahi Z, Gupta I, et al. The Not5 subunit of the ccr4-not complex connects transcription and translation. PLoS Genet. 2014;10(10):e1004569.

48. Gupta I, Villanyi Z, Kassem S, Hughes C, Panasenko OO, Steinmetz LM, et al. Translational Capacity of a Cell Is Determined during Transcription Elongation via the Ccr4-Not Complex. Cell Rep. 2016;15(8):1782–94.

49. Kassem S, Villanyi Z, Collart MA. Not5-dependent co-translational assembly of Ada2 and Spt20 is essential for functional integrity of SAGA. Nucleic Acids Res. 2017;45(3):1186–99.

50. Buschauer R, Matsuo Y, Sugiyama T, Chen YH, Alhusaini N, Sweet T, et al. The Ccr4-Not complex monitors the translating ribosome for codon optimality. Science. 2020;368(6488).

51. Benne R, Hershey JW. The mechanism of action of protein synthesis initiation factors from rabbit reticulocytes. J Biol Chem. 1978;253(9):3078–87.

52. Schuller AP, Wu CC, Dever TE, Buskirk AR, Green R. eIF5A Functions Globally in Translation Elongation and Termination. Mol Cell. 2017;66(2):194–205 e5.

53. Halter D, Collart MA, Panasenko OO. The Not4 E3 ligase and CCR4 deadenylase play distinct roles in protein quality control. PLoS One. 2014;9(1):e86218.

54. Garcia M, Delaveau T, Goussard S, Jacq C. Mitochondrial presequence and open reading frame mediate asymmetric localization of messenger RNA. EMBO Rep. 2010;11(4):285–91.

55. Jan CH, Williams CC, Weissman JS. Principles of ER cotranslational translocation revealed by proximity-specific ribosome profiling. Science. 2014;346(6210):1257521.

56. Shurtleff MJ, Itzhak DN, Hussmann JA, Schirle Oakdale NT, Costa EA, Jonikas M, et al. The ER membrane protein complex interacts cotranslationally to enable biogenesis of multipass membrane proteins. Elife. 2018;7.

57. Williams CC, Jan CH, Weissman JS. Targeting and plasticity of mitochondrial proteins revealed by proximity-specific ribosome profiling. Science. 2014;346(6210):748–51.

58. Vardi-Oknin D, Arava Y. Characterization of Factors Involved in Localized Translation Near Mitochondria by Ribosome-Proximity Labeling. Front Cell Dev Biol. 2019;7:305.

59. Garcia-Rodriguez LJ, Gay AC, Pon LA. Puf3p, a Pumilio family RNA binding protein, localizes to mitochondria and regulates mitochondrial biogenesis and motility in budding yeast. J Cell Biol. 2007;176(2):197–207.

60. Zhang Y, Pelechano V. Application of high-throughput 5’P sequencing for the study of co-translational mRNA decay. STAR Protoc. 2021;2(2):100447.

61. Ingolia NT, Lareau LF, Weissman JS. Ribosome profiling of mouse embryonic stem cells reveals the complexity and dynamics of mammalian proteomes. Cell. 2011;147(4):789–802.

62. Hafner M, Landthaler M, Burger L, Khorshid M, Hausser J, Berninger P, et al. Transcriptome-wide identification of RNA-binding protein and microRNA target sites by PAR-CLIP. Cell. 2010;141(1):129–41.

63. Schaughency P, Merran J, Corden JL. Genome-wide mapping of yeast RNA polymerase II termination. PLoS Genet. 2014;10(10):e1004632.

64. Cooper KF, Scarnati MS, Krasley E, Mallory MJ, Jin C, Law MJ, et al. Oxidative-stress-induced nuclear to cytoplasmic relocalization is required for Not4-dependent cyclin C destruction. J Cell Sci. 2012;125(Pt 4):1015–26.

65. Panasenko OO, Collart MA. Not4 E3 ligase contributes to proteasome assembly and functional integrity in part through Ecm29. Molecular and cellular biology. 2011;31(8):1610–23.

66. Webster MW, Chen YH, Stowell JAW, Alhusaini N, Sweet T, Graveley BR, et al. mRNA Deadenylation Is Coupled to Translation Rates by the Differential Activities of Ccr4-Not Nucleases. Mol Cell. 2018;70(6):1089–100 e8.

67. Ikeuchi K, Izawa T, Inada T. Recent Progress on the Molecular Mechanism of Quality Controls Induced by Ribosome Stalling. Front Genet. 2018;9:743.

68. Ikeuchi K, Tesina P, Matsuo Y, Sugiyama T, Cheng J, Saeki Y, et al. Collided ribosomes form a unique structural interface to induce Hel2-driven quality control pathways. EMBO J. 2019;38(5).

69. Gillen SL, Giacomelli C, Hodge K, Zanivan S, Bushell M, Wilczynska A. Differential regulation of mRNA fate by the human Ccr4-Not complex is driven by coding sequence composition and mRNA localization. Genome Biol. 2021;22(1):284.

70. Mugler CF, Hondele M, Heinrich S, Sachdev R, Vallotton P, Koek AY, et al. ATPase activity of the DEAD-box protein Dhh1 controls processing body formation. Elife. 2016;5.

71. Collart MA, Oliviero S. Preparation of yeast RNA. Curr Protoc Mol Biol. 2001;Chapter 13:Unit13 2.

72. Oberholzer U, Collart MA. Characterization of NOT5 that encodes a new component of the Not protein complex. Gene. 1998;207(1):61–9.

73. Azzouz N, Panasenko OO, Deluen C, Hsieh J, Theiler G, Collart MA. Specific roles for the Ccr4-Not complex subunits in expression of the genome. Rna. 2009;15(3):377–83.

74. Zhang Y, Pelechano V. High-throughput 5’P sequencing enables the study of degradation-associated ribosome stalls. bioRxiv. 2020.

75. Tagwerker C, Zhang H, Wang X, Larsen LS, Lathrop RH, Hatfield GW, et al. HB tag modules for PCR-based gene tagging and tandem affinity purification in Saccharomyces cerevisiae. Yeast. 2006;23(8):623–32.

76. Kubik S, O’Duibhir E, de Jonge WJ, Mattarocci S, Albert B, Falcone JL, et al. Sequence-Directed Action of RSC Remodeler and General Regulatory Factors Modulates +1 Nucleosome Position to Facilitate Transcription. Mol Cell. 2018;71(1):89–102 e5.

77. Kirchner S, Cai Z, Rauscher R, Kastelic N, Anding M, Czech A, et al. Alteration of protein function by a silent polymorphism linked to tRNA abundance. PLoS Biol. 2017;15(5):e2000779.

78. Martin M. Cutadapt Removes Adapter Sequences from High-Throughput Sequencing Reads. EMBnet Journal. 2011;17:10–2.

79. Love MI, Huber W, Anders S. Moderated estimation of fold change and dispersion for RNA-seq data with DESeq2. Genome Biol. 2014;15(12):550.

80. Langmead B, Trapnell C, Pop M, Salzberg SL. Ultrafast and memory-efficient alignment of short DNA sequences to the human genome. Genome Biol. 2009;10(3):R25.

81. Comoglio F, Sievers C, Paro R. Sensitive and highly resolved identification of RNA-protein interaction sites in PAR-CLIP data. BMC Bioinformatics. 2015;16:32.

82. Langmead B, Salzberg SL. Fast gapped-read alignment with Bowtie 2. Nat Methods. 2012;9(4):357–9.

83. Wilcoxon F. Individual comparisons of grouped data by ranking methods. J Econ Entomol. 1946;39:269.

