## Supplementary Figures and legends for "Not1 and Not4 inversely determine mRNA solubility that sets the dynamics of co-translational events"

#### Supplementary Figure legends

**Figure S1. Comparison of mRNAs that are more or less soluble. A**. GO-term analysis of the most and least soluble mRNAs in wild type cells. **B**. Metagene profiles of 5'P decay intermediates for mRNAs separated in equivalent sized groups according to level of expression and length for soluble and total RNAs. **C**. Metagene analysis of 5'P decay intermediates in soluble RNA pools (left) and of ribosome footprints (17) (right) for wild type cells. **D**. Metagene analysis of 5'ends of RNA-Seq reads for total and soluble RNA pools. **E**. Heat maps (upper panels) and box plot analysis (bottom panels) of log2FC ratios of 5'P-Seq reads or 5' end of RNA-Seq reads between 70-90% of CDSs and 10-30% of CDSs for soluble and total RNA pools for all mRNAs with at least 5 reads in region 10-30% or 70-90% for both 5'P-Seq and RNA-Seq. **F**. Scatterplot comparing 5'P-RDO differences between second half of CDSs to the first half of CDSs for soluble (left) and total (right) RNA pools from wild type cells.

**Figure S2. mRNAs that are more or less soluble upon Not1 and Not4 depletion are enriched for different GO-terms. A**. Total protein extracts of cells carrying the Not4 (left) and Not5 (right) degrons were prepared before (-) and after (+) treatment with auxin for the indicated times. Western blot analysis of equivalent amounts of protein were analyzed with antibodies to (Myc, top rows), Not1 (bottom rows) and either Not5 (left middle row) or Not4 (right middle row) as indicated. **B**. Scatterplot analysis comparing changes in mRNA solubilities upon Not4 and Not5 (left panel) or Not1 and Not5 (right panel) depletions. **C**. GO-term analysis of the red, blue, green and orange mRNAs from **Figure 2C**. **D**. Relative levels (in percent) of reads of 5'P decay intermediates mapping to the combined *S. cerevisiae/S pombe* genomes in the different RNA samples that map to *S.pombe.*

**Figure S3. mRNAs whose solubilities are inversely regulated by Not1 and Not4 show different changes in co-translational decay upon Not1, Not4 and Not5 depletions**. **A.** Scatterplot analyses comparing changes in 5'P-RDOs for soluble RNAs upon Not5 depletion to RDO changes in *not5Δ* compared to wild type (17). **B**. Scatterplot analyses comparing changes in 5'P-RDOs in cells upon depletion of Not1 (*not1d*/WT), Not4 (*not4d*/WT) or Not5 (*not5d*/WT) in soluble (sol) versus total (tot) RNA pools.

**Figure S4**. **Quality control of the Not4 PAR-CLIP. A**. Scatterplot (top) and distribution of differences (bottom) comparing Not4 cross-linking in the duplicate samples. **B**. Scatterplot (top) and distribution of differences (bottom) comparing Not4 cross-linking by T to C transitions or overall reads and to coding sequences (CDS, left) and introns (right). **C**. Scatterplots comparing cross-linking of Not4 and RNAPII (63) to different sequences (CDS, 5'UTR, 3'UTR and introns). **D**. Same as for panel **C** but for antisense transcripts. **E**. Metagene profiles of Not4 and RNAPII cross-linking (63) to specific regions: around the start (left), on the ORF (middle) and around the stop (right). **F**. Scatterplot comparing overall cross-linking of Not4 and RNAPII. **G**. GO-term analysis of mRNAs more highly cross-linked by Not4 than RNAPII compared to the mean relative cross-linking of Not4 to RNAPII. **H-J**. Box plot analysis comparing the PAR-CLIP of Not4 (**H**), RNAPII (**I**) or RNA-Seq RPKMs (**J**) to the different categories in **Figure 2C**. Significance of differences is indicated at the top of the box plots - p-values are calculated using a two-sided Welch two sample t-test.

**Supplementary Tables**

**Table S1. 5'P-Seq and RNA-Seq of soluble and total RNA pools before and after depletion of Not1, Not4 and Not5.**

**Table S2. Changes in solubility upon Not1, Not4 and Not5 depletion.**

**Table S3. Not4 and RNAPII PAR-CLIP**.

**Table S4**. **Not1-ChEC-Seq**

### Figure S1

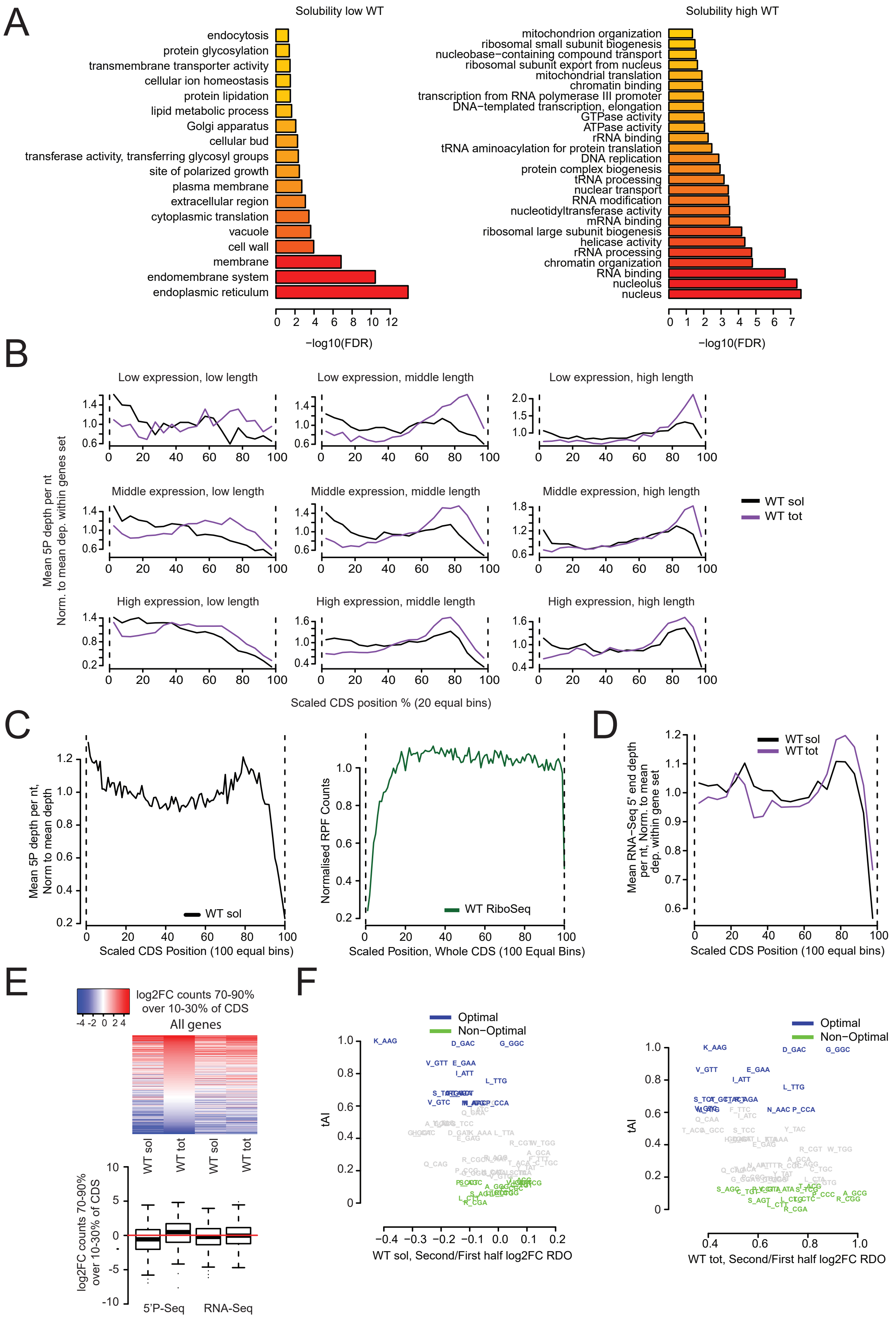

A
Solubility low WT
endocytosis
protein glycosylation
transmembrane transporter activity
cellular ion homeostasis
protein lipidation
lipid metabolic process
Golgi apparatus
cellular bud
transferase activity, transferring glycosyl groups
site of polarized growth
plasma membrane
extracellular region
cytoplasmic translation
vacuole
cell wall
membrane
endomembrane system
endoplasmic reticulum
−log10(FDR)
Solubility high WT
mitochondrion organization
ribosomal small subunit biogenesis
nucleobase−containing compound transport
ribosomal subunit export from nucleus
mitochondrial translation
chromatin binding
transcription from RNA polymerase III promoter
DNA−templated transcription, elongation
GTPase activity
ATPase activity
rRNA binding
tRNA aminoacylation for protein translation
DNA replication
protein complex biogenesis
tRNA processing
nuclear transport
RNA modification
nucleotidyltransferase activity
mRNA binding
ribosomal large subunit biogenesis
helicase activity
rRNA processing
chromatin organization
RNA binding
nucleolus
nucleus
−log10(FDR)
B
Low expression, low length
Low expression, middle length
Low expression, high length
Middle expression, low length
Middle expression, middle length
Middle expression, high length
High expression, low length
High expression, middle length
High expression, high length
Mean 5P depth per nt
Norm. to mean dep. within genes set
WT sol
WT tot
Scaled CDS position % (20 equal bins)
C
Mean 5P depth per nt, Norm to mean depth
WT sol
Scaled CDS Position (100 equal bins)
Normalised RPF Counts
WT RiboSeq
Scaled Position, Whole CDS (100 Equal Bins)
D
Mean RNA−Seq 5' end depth per nt, Norm. to mean dep. within gene set
WT sol
WT tot
Scaled CDS Position (100 equal bins)
E
log2FC counts 70-90% over 10-30% of CDS
All genes
WT sol
WT tot
WT sol
WT tot
log2FC counts 70-90% over 10-30% of CDS
5'P-Seq
RNA-Seq
F
Optimal
Non−Optimal
tAI
WT sol, Second/First half log2FC RDO
WT tot, Second/First half log2FC RDO

### Figure S2

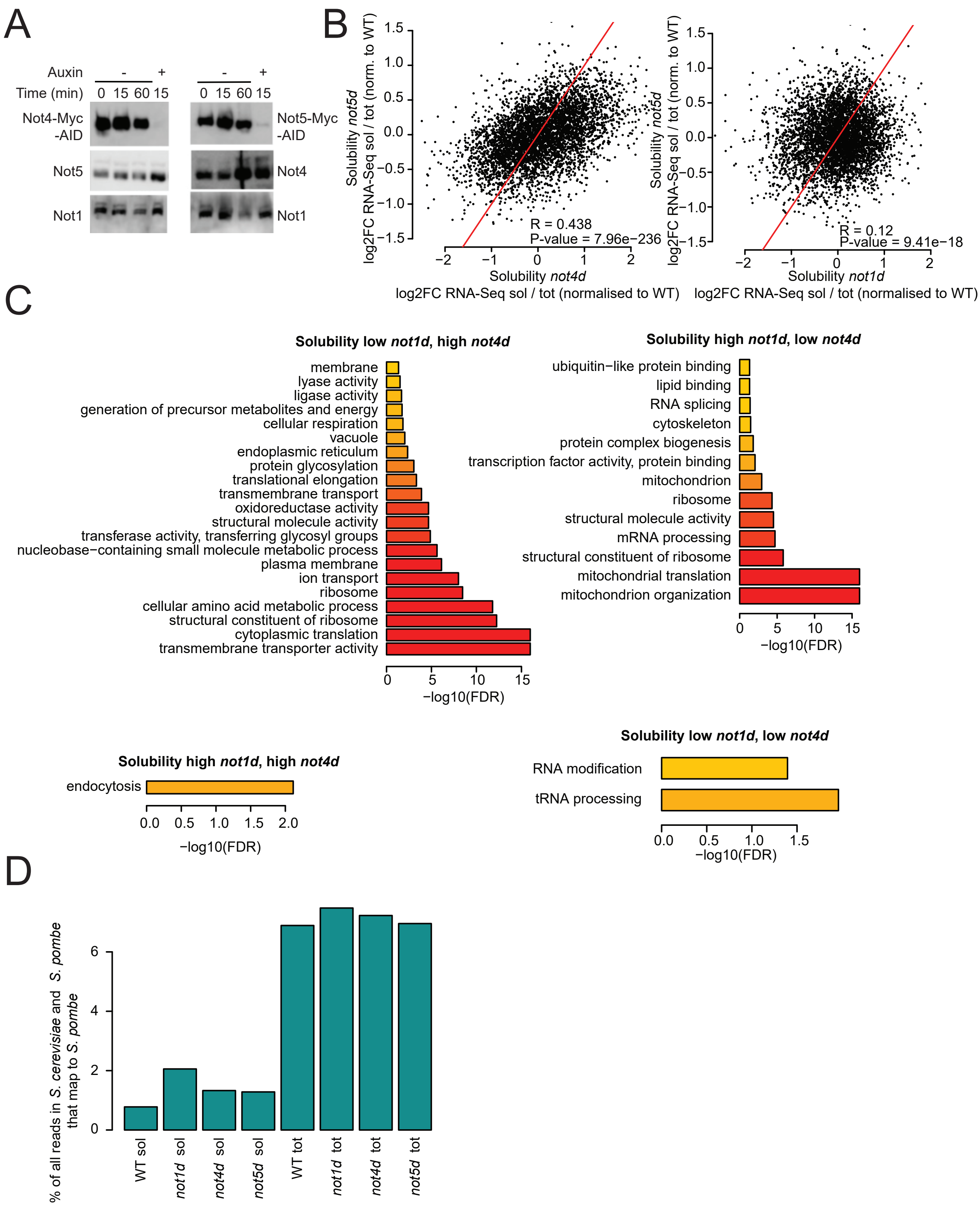

A
Auxin
-
+
Time (min)
0 15 60 15
Not4-Myc -AID
Not5
Not1
Not5-Myc -AID
Not4
Not1
B
Solubility not5d
log2FC RNA-Seq sol / tot (norm. to WT)
R = 0.438
P-value = 7.96e−236
Solubility not4d
log2FC RNA-Seq sol / tot (normalised to WT)
R = 0.12
P-value = 9.41e−18
Solubility not1d
log2FC RNA-Seq sol / tot (normalised to WT)
C
Solubility low not1d, high not4d
membrane
lyase activity
ligase activity
generation of precursor metabolites and energy
cellular respiration
vacuole
endoplasmic reticulum
protein glycosylation
translational elongation
transmembrane transport
oxidoreductase activity
structural molecule activity
transferase activity, transferring glycosyl groups
nucleobase−containing small molecule metabolic process
plasma membrane
ion transport
ribosome
cellular amino acid metabolic process
structural constituent of ribosome
cytoplasmic translation
transmembrane transporter activity
−log10(FDR)
Solubility high not1d, low not4d
ubiquitin−like protein binding
lipid binding
RNA splicing
cytoskeleton
protein complex biogenesis
transcription factor activity, protein binding
mitochondrion
ribosome
structural molecule activity
mRNA processing
structural constituent of ribosome
mitochondrial translation
mitochondrion organization
−log10(FDR)
Solubility high not1d, high not4d
endocytosis
−log10(FDR)
Solubility low not1d, low not4d
RNA modification
tRNA processing
−log10(FDR)
D
% of all reads in S. cerevisiae and S. pombe that map to S. pombe
WT sol
not1d sol
not4d sol
not5d sol
WT tot
not1d tot
not4d tot
not5d tot

### Figure S3

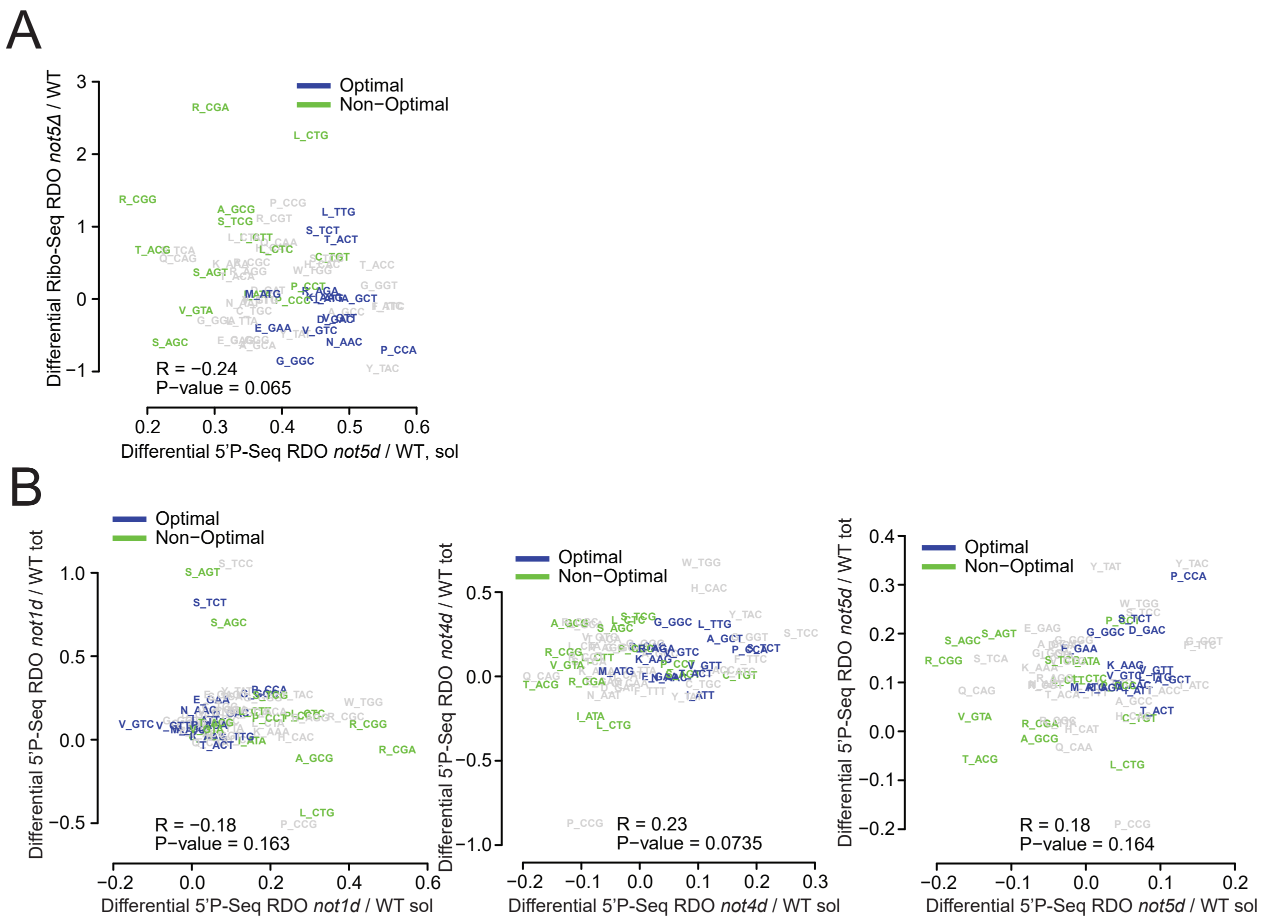

A
Optimal
Non−Optimal
Differential Ribo-Seq RDO not5Δ / WT
Differential 5'P-Seq RDO not5d / WT, sol
R = −0.24
P−value = 0.065
B
Optimal
Non−Optimal
Differential 5'P-Seq RDO not1d / WT tot
Differential 5'P-Seq RDO not1d / WT sol
R = −0.18
P−value = 0.163
Optimal
Non−Optimal
Differential 5'P-Seq RDO not4d / WT tot
Differential 5'P-Seq RDO not4d / WT sol
R = 0.23
P−value = 0.0735
Optimal
Non−Optimal
Differential 5'P-Seq RDO not5d / WT tot
Differential 5'P-Seq RDO not5d / WT sol
R = 0.18
P−value = 0.164

Figure S4

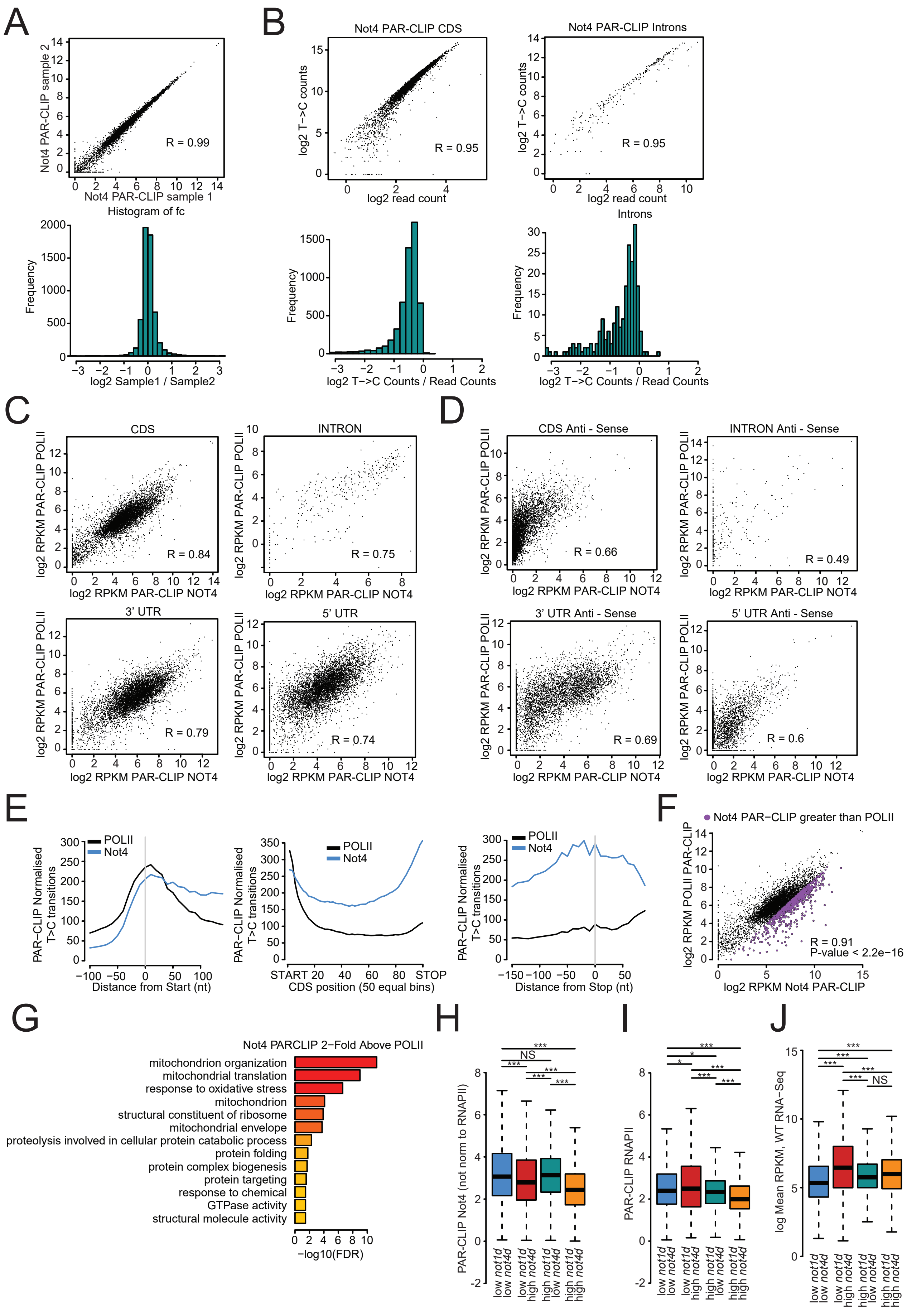
A
Not4 PAR-CLIP sample 2
Not4 PAR-CLIP sample 1
R = 0.99
Histogram of fc
Frequency
log2 Sample1 / Sample2
B
Not4 PAR-CLIP CDS
log2 T->C counts
log2 read count
R = 0.95
Not4 PAR-CLIP Introns
R = 0.95
Introns
log2 T−>C Counts / Read Counts
C
CDS
R = 0.84
INTRON
R = 0.75
3' UTR
R = 0.79
5' UTR
R = 0.74
log2 RPKM PAR-CLIP POLII
log2 RPKM PAR-CLIP NOT4
D
CDS Anti - Sense
R = 0.66
INTRON Anti - Sense
R = 0.49
3' UTR Anti - Sense
R = 0.69
5' UTR Anti - Sense
R = 0.6
E
POLII
Not4
PAR−CLIP Normalised T>C transitions
Distance from Start (nt)
CDS position (50 equal bins)
START
STOP
Distance from Stop (nt)
F
Not4 PAR−CLIP greater than POLII
log2 RPKM POLII PAR-CLIP
log2 RPKM Not4 PAR-CLIP
R = 0.91
P-value < 2.2e−16
G
Not4 PARCLIP 2−Fold Above POLII
mitochondrion organization
mitochondrial translation
response to oxidative stress
mitochondrion
structural constituent of ribosome
mitochondrial envelope
proteolysis involved in cellular protein catabolic process
protein folding
protein complex biogenesis
protein targeting
response to chemical
GTPase activity
structural molecule activity
−log10(FDR)
H
PAR-CLIP Not4 (not norm to RNAPII)
NS
I
PAR-CLIP RNAPII
J
log Mean RPKM, WT RNA−Seq
NS
low not1d low not4d
low not1d high not4d
high not1d low not4d
high not1d high not4d
